# Identification of Cross-Reactive CD8^+^ T Cell Receptors with High Functional Avidity to a SARS-CoV-2 Immunodominant Epitope and Its Natural Mutant Variants

**DOI:** 10.1101/2020.11.02.364729

**Authors:** Chao Hu, Meiying Shen, Xiaojian Han, Qian Chen, Luo Li, Siyin Chen, Jing Zhang, Fengxia Gao, Wang Wang, Yingming Wang, Tingting Li, Shenglong Li, Jingjing Huang, Jianwei Wang, Ju Zhu, Dan Chen, Qingchen Wu, Kun Tao, Da Pang, Aishun Jin

**Affiliations:** Department of Immunology, College of Basic Medicine, Chongqing Medical University, 1 Yixueyuan Road, Yuzhong District, Chongqing, 400016, China; Chongqing Key Laboratory of Cancer Immunology Translational Medicine, Chongqing Medical University, 1 Yixueyuan Road, Yuzhong District, Chongqing, 400016, China; Department of Breast Surgery, Harbin Medical University Cancer Hospital, 150 Haping Road, Nangang District, Harbin, 150081, China; Department of Cardiothoracic Surgery, The First Affiliated Hospital of Chongqing Medical University, 1 Youyi Road, Yuzhong District, Chongqing, 400016, China

**Keywords:** SARS-CoV-2, CD8^+^ T cell, T cell epitope, TCR, Lung organoid, HLA class I

## Abstract

Despite the growing knowledge of T cell responses and their epitopes in COVID-19 patients, there is a lack of detailed characterizations for T cell-antigen interactions and T cell functions. Using a peptide library predicted with HLA class I-restriction, specific CD8^+^ T cell responses were identified in over 75% of COVID-19 convalescent patients. Among the 15 SARS-CoV-2 epitopes identified from the S and N proteins, N^361-369^ (KTFPPTEPK) was the most dominant epitope. Importantly, we discovered 2 N^361-369^-specific T cell receptors (TCRs) with high functional avidity, and they exhibited complementary cross-reactivity to reported N^361-369^ mutant variants. In dendritic cells (DCs) and the lung organoid model, we found that the N^361-369^ epitope could be processed and endogenously presented to elicit the activation and cytotoxicity of CD8^+^ T cells *ex vivo*. Our study evidenced potential mechanisms of cellular immunity to SARS-CoV-2, illuminating natural ways of viral clearance with high relevancy in the vaccine development.

## INTRODUCTION

Severe acute respiratory syndrome coronavirus 2 (SARS-CoV-2) results in the global pandemic of COVID-19 (Dong et al., 2020; Zhou et al., 2020). So far, over 40 million positive cases of SARS-CoV-2 infection have been reported with one million deaths worldwide (*https://covid19.who.int/*). Despite the fact that most patients can recover spontaneously with supportive care, COVID-19 mortality rate is still a worrying social concern. A better understanding of the adaptive immunity to SARS-CoV-2 infection is important to assist vaccine development and evaluation.

SARS-CoV-2 contains four structural proteins, namely nucleoprotein (N), membrane protein (M), envelope protein (E) and spike glycoprotein (S) (Zhou et al., 2020). SARS-CoV-2 S protein is responsible for the virus-target cell recognition (Hoffmann et al., 2020; Walls et al., 2020; Zhou et al., 2020). It has been demonstrated that the S protein was capable of inducing not only humoral immunity but also cellular immunity (Cao et al., 2020; Chen et al., 2020; Grifoni et al., 2020b; Ju et al., 2020; Le Bert et al., 2020a; Ni et al., 2020; Shi et al., 2020; Weiskopf et al., 2020). And the S protein has always been the primary consideration for SARS-CoV-2 vaccine development (Corbett et al., 2020a; Mercado et al., 2020b; Yu et al., 2020). Besides the S protein, the N protein can also elicit high levels of T cell responses in COVID-19 patients, highlighting the possibility that S and N are both predominant targets for T cell responses in SARS-CoV-2 (Grifoni et al., 2020b; Le Bert et al., 2020b; Peng et al., 2020). It is still unknown whether SARS-CoV-2 is effective to induce long-lasting protective immunity due to the limited time window post-outbreak.

Pandemic outbreaks of infectious diseases resulted from pathogenic human coronaviruses, such as SARS-CoV and middle eastern respiratory syndrome coronavirus (MERS-CoV), have caused high mortality in the past two decades. For SARS-CoV, evidences supporting T cell immunity to its N protein has been reported, with some still observed over 17 years later in SARS-CoV infected individuals (Le Bert et al., 2020b). Because SARS-CoV N protein exhibits more than 90% homology of amino acid sequence to that of SARS-CoV-2 (Grifoni et al., 2020a), it is natural to suspect that T cell responses to the N protein of SARS-CoV-2 may also provide cellular immune protection, which may last for a long time. Recently, using predicted or established peptide libraries of the S and N proteins, a number of SARS-CoV-2 T cell epitopes were identified in COVID-19 convalescent patients across independent studies, providing important clues for vaccine development (Chour et al., 2020; Ferretti et al., 2020; Le Bert et al., 2020b; Nelde et al., 2020; Peng et al., 2020; Poran et al., 2020; Shomuradova et al., 2020; Snyder et al., 2020). However, detailed information of how SARS-CoV-2 induces cellular immune responses is currently limited.

Cellular immunity against viral infections is mediated by cytotoxic T cells, which eliminate the viruses by killing infected cells. To trigger optimal T cells responses, viral antigens must be processed before binding to the major histocompatibility complex (MHC) molecules to form the peptide-MHC complex (pMHC), which is then presented on the cell membrane. When TCRs identify the cognate pMHC, together with matching CD8 co-receptor binding to the MHC, the T cells will be activated. Such activation is dependent on the binding kinetics of the TCR-pMHC, which is influenced by the density of processed epitopes on the membrane of the virus-infected cells or antigen presenting cells (APC) (Campillo-Davo et al., 2020; Gonzalez et al., 2005). Particularly, CD8 co-receptor is indispensable for enhanced activation of T cells with low-affinity TCRs, but not for T cells with high-affinity TCRs. (Campillo-Davo et al., 2020; Lyons et al., 2006; Tan et al., 2017). Activation of T cells independent of the CD8 co-receptor reflects the high functional avidity of their TCRs. Hence, the functional avidity of TCRs is a critical parameter to be evaluated for determining the pMHC recognition and posterior T-cell activation, which is essential for the killing of virus-infected cells and effective viral clearance (Campillo-Davo et al., 2020).On the other hand, RNA viruses have a tendency to accumulate mutations over time (Duffy, 2018; Villa et al., 2020), which present challenges for T cell-based vaccine development. Therefore, it is important to identify predominant epitopes together with diverse TCRs carrying broader cross-reactivity to SARS-CoV-2 and its mutant variants, which can provide detailed information to understand the mechanisms of T cell-mediated viral clearance in COVID-19 patients.

In this study, we evaluated SARS-CoV-2 specific CD8^+^ T cell responses in COVID-19 convalescent patients with HLA-A2, HLA-A24 or HLA-A11 allele, the 3 most prominent HLA-A alleles in the Asian population (He et al., 2018). A total of 15 epitopes derived from SARS-CoV-2 S and N were identified. The most dominant epitope N^361-369^ (KTFPPTEPK) were characterized in detail, including the wild type and clinically presented mutant variants of N^361-369^ and the functional avidity of N^361-369^-specific TCRs. Our study is, to our knowledge, the first to reveal that an immunodominant CD8^+^ T cell epitope derived from SARS-CoV-2 N can be intracellularly processed and presented by DCs and lung organoids, leading to robust T cell activation and cytotoxicity. Our findings illustrated potential mechanisms of CD8^+^ T cell immunity against SARS-CoV-2 infection, which could be of great use to understand the viral control process and to facilitate the vaccine development for COVID-19.

## RESULTS

### Identification of SARS-CoV-2 Specific CD8^+^ T Cell Epitopes

It has been reported that SARS-CoV-2 S and N proteins were the predominant T cell targets in COVID-19 patients (Grifoni et al., 2020b). In order to predict SARS-CoV-2 specific CD8^+^ T cell epitopes, the NetMHCpan 4.0 algorithm was used to analyse the binding of 9-mer peptides derived from the full lengths of SARS-CoV-2 S and N proteins, with HLA-A*02:01, HLA-A*24:02 and HLA-A*11:01, the 3 most frequent HLA class I A alleles in the Asian population. We synthesized a peptide library of 72 predicted SARS-CoV-2 epitopes, which overlapped 29 SARS-CoV epitopes, together with additional 6 confirmed SARS-CoV epitopes (Table S1) (Ahmed et al., 2020; Grifoni et al., 2020a). Among these 78 peptides, 62 were from the S protein and 16 were from the N protein, with 3 different HLA-A restrictions (Table S2).

In parallel, we collected a total of 37 COVID-19 convalescent PBMC samples and subjected them for the HLA typing analysis. Among the 20 HLA-A2^+^ samples and the 12 HLA-A24^+^ samples revealed by flow cytometric assay, we selected 8 samples from each group. A matching 8 samples with HLA-A*11:01^+^ genotype was identified by RT-PCR, which included 2 overlapping with the HLA-A2^+^ group and 2 with the HLA-A24^+^ group (Table S3). These PBMCs were stimulated for 10 days in the presence of individual peptide mixtures and IL-2, and the induction of IFN-γ secretion was determined by the enzyme-linked immunospot (ELISPOT)assay. We found that 75% COVID-19 convalescent PBMCs demonstrated positive T cell responses when stimulated by corresponding peptide mixtures. Particularly, the HLA-A2^+^ group showed dominant responses to Mixture (M) 1/3/5, the HLA-A24^+^ group to M6/7 and the HLA-A*11:01^+^ group to M14 (Figures 1A-1C and S1). Next, individual peptides from each positive mixture were applied to identify specific T cell epitopes, and 15 SARS-CoV-2 specific peptides were found to be recognized by the convalescent PBMCs (Figures 1A-1C, S2 and S3; Tables 1 and S5). Among the 15 positive peptides, 5 peptides were derived from the N protein, and the rest were derived from the S protein (Table 1). Four dominant peptides were identified, with at least one peptide from each HLA group (Table 1). Two of these peptides, P63 (N^338-346^) and P77 (N^219-227^), were both HLA-A*02:01-restricted, and they could induce IFN-γ release in 4/8 and 2/8 samples, respectively (Figures 1A and S2; Table 1). The HLA-A*24:02-restricted P45 (S^448-456^) was located at the receptor binding domain of the S protein and could be recognized by 2/8 samples tested (Figure 1B; Table 1). Importantly, 5 out of 8 samples showed positive responses to the HLA-A*11:01-restricted P64 (N^361-369^). Hence we termed P64, or N^361-369^, as the most dominant epitope in this study (Figure 1C; Table 1). Of note, IFN-γ induction was not observed when these 4 dominant peptides were used to stimulate PBMCs of healthy individuals (Figure S4; Table S4). Overall, these data demonstrated that 15 predicted peptides derived from the S and the N protein sequences could induce SARS-CoV-2-specific CD8^+^ T cell responses in about 75% of COVID-19 convalescent patient samples, and at least 4 peptides were identified to be dominant CD8^+^ T cell epitopes.

**Figure 1.**
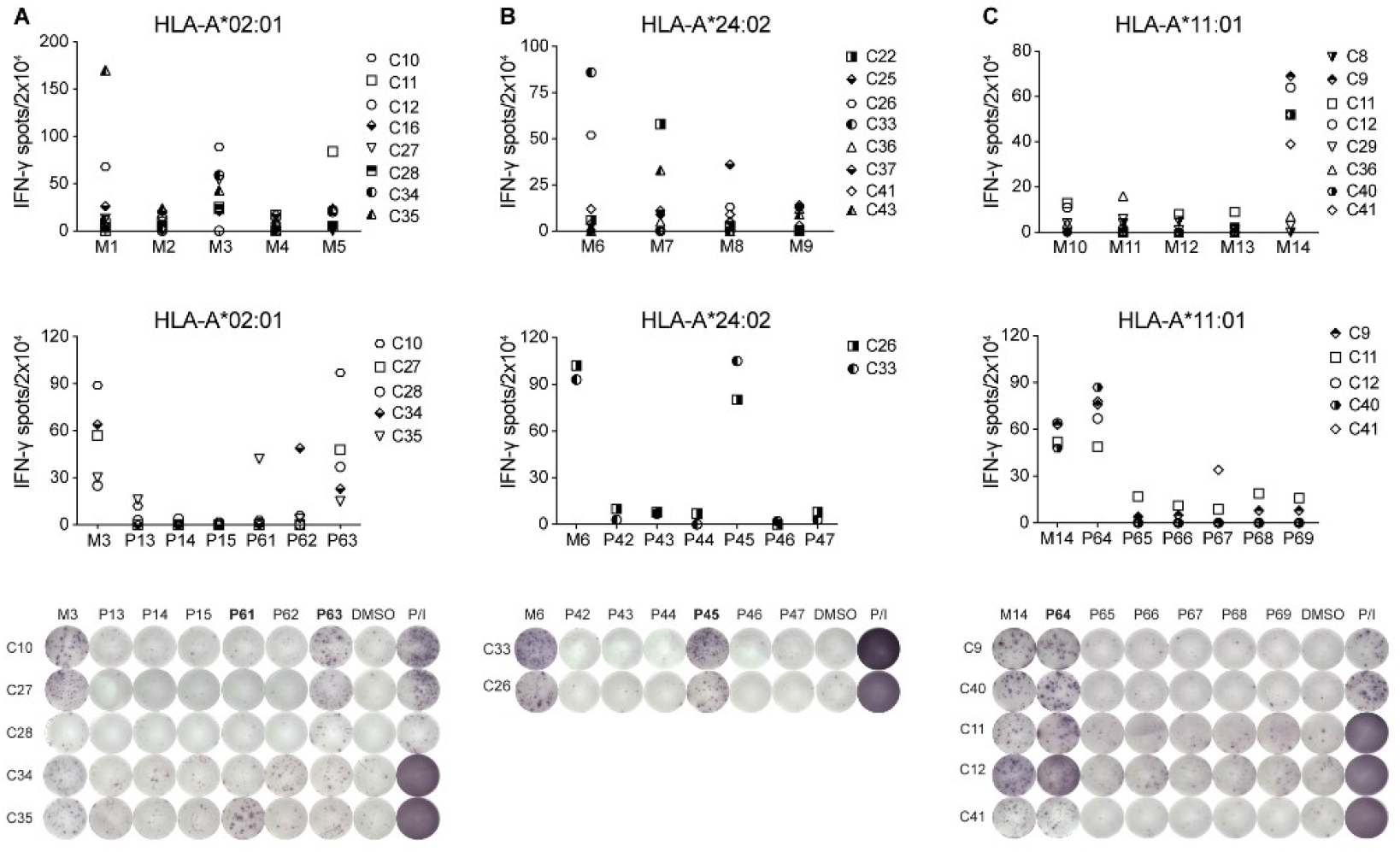
CD8^+^ T Cell Responses of SARS-CoV-2 Peptides. IFN-γ ELISPOT assay results for the PBMCs of COVID-19 convalescent patients with (A) HLA-A2^+^, (B) HLA-A24^+^ or (C) HLA-A*11:01^+^ allele, stimulated with mixed peptides (top) and single reactive peptide (middle and bottom). A stimulation with an equimolar amount of DMSO was performed as negative control, and Phorbol 12-myristate 13-acetate (PMA)/Ionomycin (P/I) was performed as positive control. Results are expressed as number of peptide-specific IFN-γ spots/2×10^4^ PBMCs of each patient, substracted those from the corresponding DMSO groups. N = 8. For peptide grouping: 26 HLA-A*02:01-restricted peptides were pooled into 5 mixtures (M1-M5), 22 HLA-A*24:02-restricted peptides were pooled into 4 mixtures (M6-M9), and 30 HLA-A*11:01-restricted peptides were pooled into 5 mixtures (M10-M14). Each mixture contained 3-6 peptides. Positive peptide mixture and single peptide with the most dominant reactivity were marked in bold. For the single dominant peptide, P63 was from M3, P45 was from M6 and P64 was from M14. C8-C41: COVID-19 convalescent patients. Data were respresentative of two independent experiments.

**Table 1.**
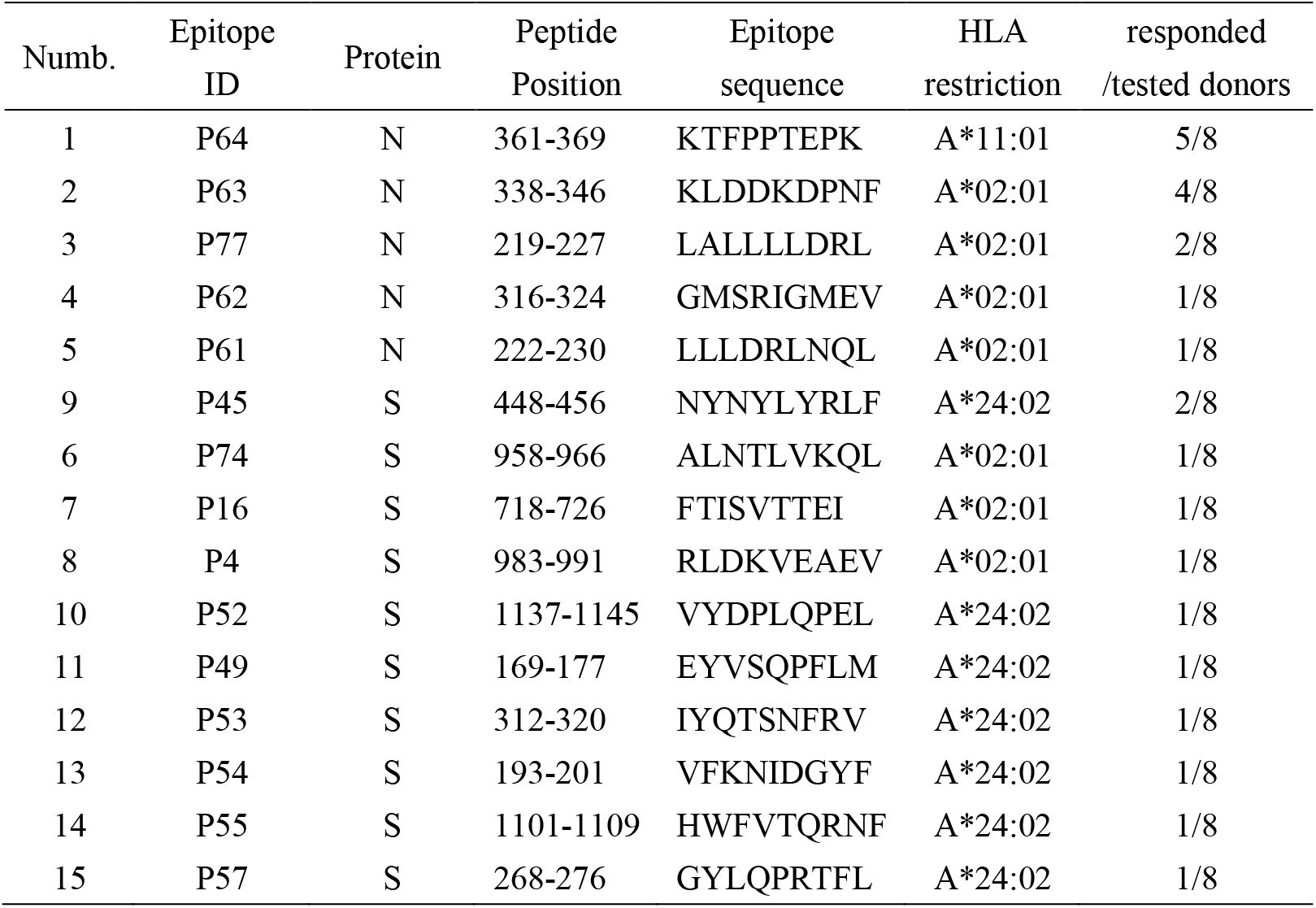
Identification of CD8^+^ T Cell Epitopes of SARS-CoV-2 in COVID-19 Convalescent Patients.

### Identification of Specific CD8^+^ TCRs to SARS-CoV-2 Dominant Epitopes

To identify the dominant epitope-specific TCRs, we selected P64, P45 and P63, the most dominant epitopes from the 3 HLA-A alleles. After stimulated with these epitopes, the populations of IFN-γ secreting T cells from the corresponding COVID-19 convalescent patient samples were sorted at single cell levels (Figure 2A). The TCRs of sorted cells were amplified (Hamana et al., 2016), and the sequences of the variable regions of α- and β-chains were analysed. In the P64-specific TCR repertoire, we found 7 dominant TCR clones (TCR 1-7), with the most dominant clone 1 (TCR 1) accounted for 17.82% (Figure 2B). Similarly, 3 dominant P63 TCR clones and 5 dominant P45 TCR clones were obtained, with the most dominant clone accounted for 10.34% and 33.34% respectively (Figure 2B). These results showed that COVID-19 convalescent patients with these three HLA-A restrictions could produce diverse T cell clones for the recognition of corresponding dominant epitopes.

**Figure 2.**
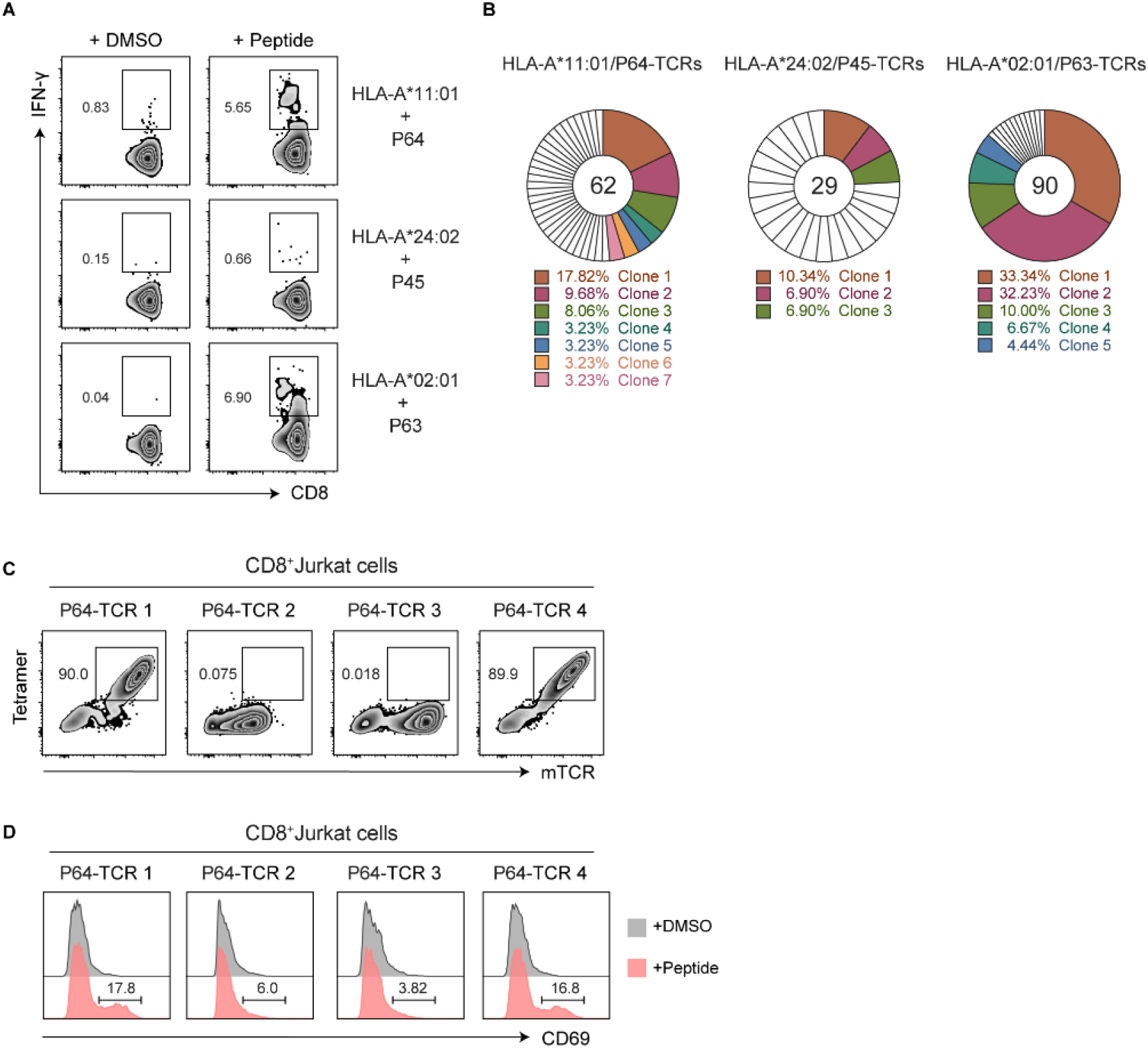
The Isolation and Validation of the SARS-CoV-2 Specific CD8^+^ TCRs. (A) After stimulated by peptide mixtures and expanded for 10 days, single IFN-γ secreting T cell from three COVID-19 convalescent PBMC samples (C40: HLA-A*11:01^+^, C33: HLA-A24^+^ and C27: HLA-A2^+^) were individually stimulated by P64, P45 and P63, from three corresponding HLA subtypes, and isolated by flow cytometric sorter. (B) The TCRs of sorted cells were amplified by RT-PCR and the TCR repertoire was analyzed with the IMGT/V-Quest tool (http://www.imgt.org/) to obtain dominant TCR clones. Unique clones were marked in white. Different colors represented relative dominant clones (copy≧2). (C-D) Specificity verificaiton for the top 4 P64-TCR clones in CD8^+^ Jurkat cells. P64-HLA-A*11:01 tetramer (Tetramer) and anti-mouse TCR (mTCR) antibody were applied to evaluate the specific binding of P64-TCRs (C). CD69 was used to determine the specific activation of TCR-transduced CD8^+^ Jurkat cells, after co-cultured with HLA-A*11:01 expressing COS-7 cells pulsed with P64 (D). Grey represented DMSO-stimulated samples and pink represented P64-stimulated samples. Data were respresentative of two independent experiments.

### Furthermore, the specificities of the P64 reactive TCRs were verified in CD8^+^

Jurkat cells, which were transduced with the top 4 dominant TCR clones (TCR 1-4). Using the P64-HLA-A*11:01 tetramer, we confirmed that only TCR 1 and TCR 4 were P64-specific (Figure 2C; Table S6). Also, with co-culture of the HLA-A*11:01-expressing COS-7 cells loaded with P64, TCR 1- or TCR 4-transduced CD8^+^ Jurkat cells could be activated, indicated by elevated CD69 expression (Figure 2D). These observations were not found with TCR 2 or TCR 3 (Figures 2C and 2D). Altogether, we successfully identified 2 different CD8^+^ TCRs targeting the SARS-CoV-2 dominant epitope P64 (N^361-369^) from the COVID-19 convalescent PBMCs.

### Characterization of the Functional Avidity of the N^361-369^-Specific TCRs

To further evaluate the functional activities of the N^361-369^-specific TCRs, TCR 1 and TCR 4 were individually transduced into the primary CD3^+^ T cells isolated from the PBMCs of healthy individuals. These cells were expanded for 12 days following the rapid expansion protocol (REP) (Jin et al., 2012), before the evaluation of their transfection efficiency by N^361-369^-HLA-A*11:01 tetramer. We found that 9.86% CD8^+^ T cells expressed TCR 1 and 6.34% expressed TCR 4 (Figure 3A). Unexpectedly, we observed higher percentages of TCR-transduced CD4^+^ T cells, with 38.4% for TCR 1 and 11.7% for TCR 4 (Figure 3A). These results indicated that TCR-transduced CD4^+^ T cells were able to recognize the N^361-369^-HLA complex with the absence of CD8 co-receptor. Also, the similar fluorescence intensities detected in the CD4^+^ T cells and in the CD8^+^ T cells implied that TCR 1 and TCR 4 might both have high avidity for the N^361-369^-HLA-A*11:01 complex (Figure 3A). To confirm the binding of these TCRs to N^361-369^-HLA-A*11:01 was independent of CD8 co-receptor, TCR 1 and TCR 4 were separately transduced to CD8^-^ and CD8^+^ Jurkat cells. Both TCR 1- and TCR 4-transduced CD8^-^ Jurkat cells showed positive tetramer staining, and comparable fluorescent signal levels were found between CD8^-^ and CD8^+^ Jurkat cells (Figure 3B). These results were in line with those of the CD4^+^ and CD8^+^ T cells (Figures 3A and 3B).

**Figure 3.**
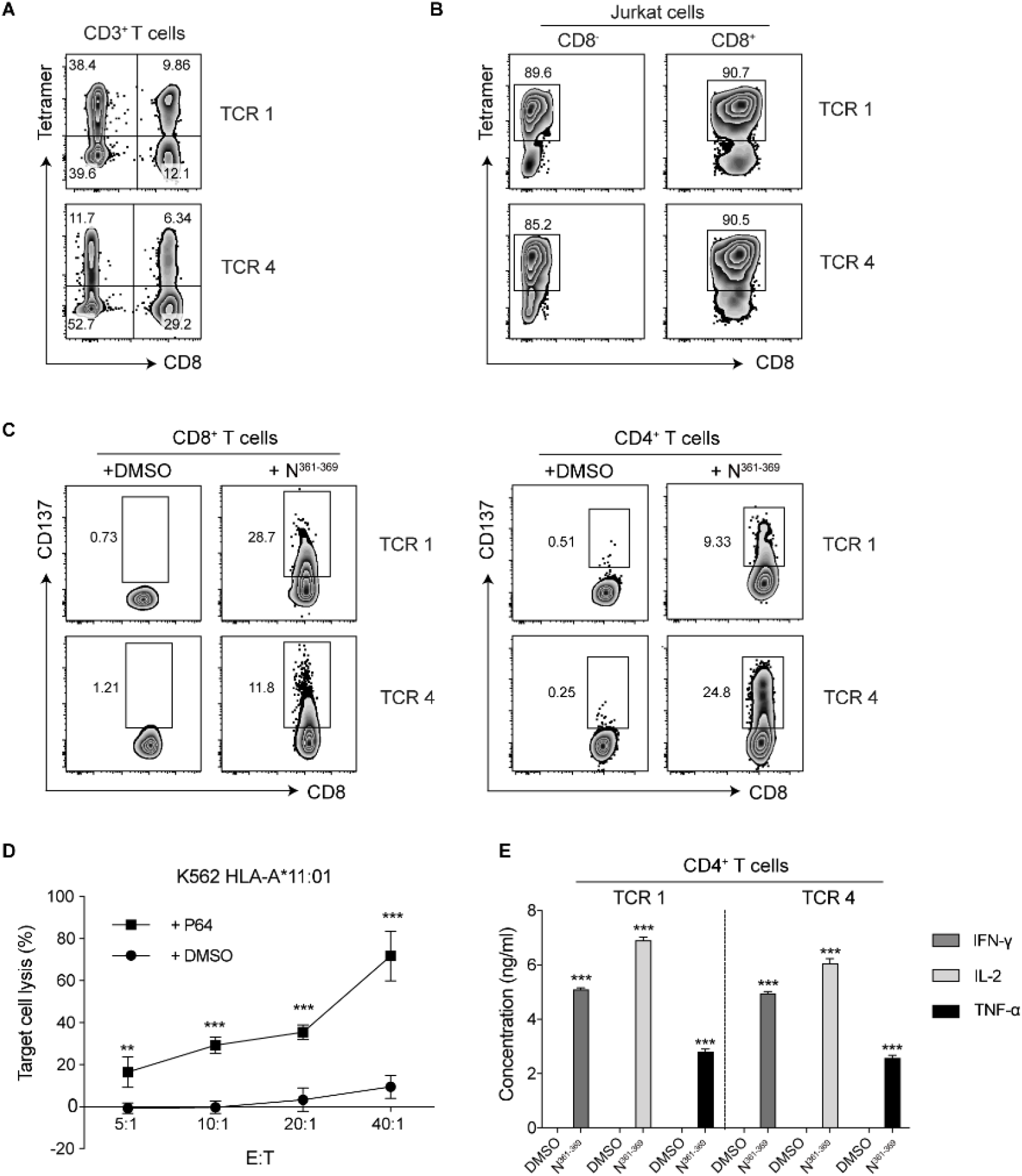
Functional Characterization of the N361-369-Specific TCR 1 and TCR 4. (A) Flow cytometric analysis of the transfection efficiency of TCR 1 and TCR 4 in CD4^+^ and CD8^+^ T cells, by N^361-369^-HLA-A*11:01 tetramer. (B) Binding assay of TCR-transduced CD8^-^ and CD8^+^ Jurkat cells with N^361-369^-HLA-A*11:01 tetramer. (C) Flow cytometric analysis of T cell activation evaluated by CD137 expression. TCR-transduced CD8^+^ T cells and CD4^+^ T cells were individually co-cultured with HLA-A*11^+^ K-562 cells pulsed with N^361-369^, or an equimolar amount of DMSO, for 24 hours. Expression of CD137 on CD8^+^ T cells (left) and CD4^+^ T cells (right) was evaluated. (D) N^361-369^ loaded HLA-A*11:01^+^ K-562 cells lysis by TCR 1-CD8^+^ T cells after 8 h at different E:T ratios. DMSO was used as negative control. Reported values are the mean of triplicates, with error bars indicating. E, Effector T cells. T, Target cells. (E) Detection of IFN-γ, IL-2 and TNF-α secretion from TCR 1 and TCR 4-transduced CD4^+^ T cells by ELISA, after co-cultured with HLA-A*11:01^+^ K-562 cells loaded with N^361-369^ for 24 h. Data were presented as mean ± SD, n = 3, **p < 0.01, ***p < 0.001. Respresentative data of two independent experiments were shown.

Next, we evaluated the functionality of TCR 1 and TCR 4 in CD8^+^ T cells and CD4^+^ T cells from expanded CD3^+^ T cells expressing TCR 1 or TCR 4. After co-cultured with HLA-A*11:01^+^ K-562 cells pulsed with N^361-369^, remarkable levels of T cell activation were detected in TCR 1- and TCR 4-CD8^+^ T cells, indicated by the upregulation of T cell activation marker CD137 expression, compared to the DMSO controls (Figure 3C). Similar results were obtained in TCR-transduced CD4^+^ T cells stimulated with N^361-369^ (Figure 3C). Moreover, we evaluated the cytotoxicity of TCR 1- and TCR 4-transduced CD8^+^ T cells against HLA-A*11:01^+^ K-562 cells, using Calcein-AM release assay. Significant levels of target cell lysis were detected in the presence of N^361-369^, relative to the DMSO controls (Figure 3D). Increasing cytotoxic efficiency was observed at the effector to target (E:T) cell ratios of 20:1 or higher, and approximately 70% target cell lysis was achieved at E:T cell ratio of 40:1 (Figure 3D). The functional activity of TCR 1- and TCR 4-transduced CD4^+^ T cells was analysed by the production of IFN-γ, IL-2 and TNF-α via cytokine secretion assay. We found that N^361-369^ could elicit significant induction of all three cytokines in these CD4^+^ T cells, comparing to an equimolar amount of DMSO (Figure 3E). These results proved that the N^361-369^-specific TCR 1 and TCR 4 could mediate the cytotoxicity of CD8^+^ T cells and the effector cytokine secretion of CD4^+^ T cells.

### Evaluation of the Affinity and the Cross-Reactivity of the N^361-369^-Specific TCRs

We addressed the functional avidity of TCR 1 and TCR 4 to the N^361-369^-HLA complex with the peptide concentrations required to induce a half-maximal activation (EC50) per IFN-γ production by the TCR-transduced T cells. The IFN-γ release assay revealed that the EC50 concentrations of TCR 1 and TCR 4 in CD8^+^ T cells were 46.83 nM and 47.17 nM respectively, when stimulated by HLA-A*11:01^+^ K-562 cells loaded with sequential ten-fold dilutions of the N^361-369^ peptide (Figure 4A). Notably, TCR 1 and TCR 4 in CD4^+^ T cells also demonstrated high functional avidity, with EC50s of 118 nM and 380.9 nM respectively, in a co-receptor independent manner (Figure 4A). These data confirmed that TCR 1 and TCR 4 both had high affinity to the N^361-369^-HLA complex in CD8^+^ and CD4^+^ T cells.

**Figure 4.**
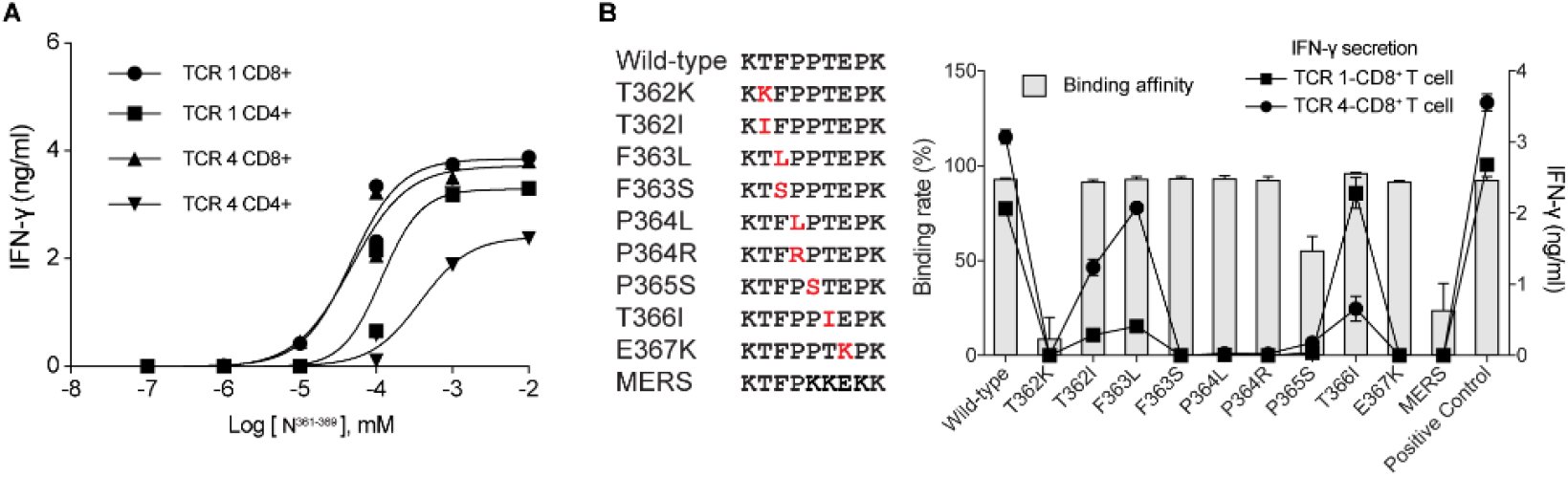
Evaluation of the Affinity and the Cross-Reactivity of TCR 1 and TCR 4 to the N361-369 Mutants. (A) EC50 analysis with IFN-γ secretion assessed by ELISA. HLA-A*11^+^ K-562 cells loaded with titrating concentrations of the N^361-369^ peptide were co-cultured with the TCR-CD4^+^ T cells or the TCR-CD8^+^ T cells for 24 h. Individual EC50 of N^361-369^ was calculated using nonlinear regression analysis. (B) Nine N^361-369^ variants were aligned with the wild type peptide and the homologous MERS peptide (left). Individual mutations were marked in red. HLA-A*11:01 binding affinity of each peptide was determined by the peptide exchange rates and shown by grey bars (n = 2). The reactivity of TCR 1- and TCR 4-CD8^+^ T cells to the N^361-369^ mutant peptides were tested by IFN-γ ELISA assay and shown in connecting lines (n = 3). Data were presented as mean + SD. Respresentative data of two independent experiments were shown.

Recently, SARS-CoV-2 mutation strains have been reported continuously. To evaluate whether TCR 1 and TCR 4 could recognize known N^361-369^ mutations, we synthesized a total of 9 mutant peptides with clinically presented point mutations of residues between 361 to 369 amino acids of the N protein (Figure 4B; Table S7) (Zhao et al., 2020). The HLA binding assay revealed that 7 different point mutations (T362I, F363L, F363S, P364L, P364R, T366I and E367K) did not affect the binding of N^361-369^ to HLA*A11:01, whereas T362K markedly diminished the ability of the N^T362K^ peptide to bind to HLA*A11:01. These results, particularly the different reactivity of N^T362I^ and N^T362K^, implied that the charge of amino acid side chain, at least, might influence the binding affinity of N^361-369^ to HLA*A11:01. We also found that the N^361-369^ homologous peptide from MERS-CoV (N^MERS^), which exhibited 3 amino acids in difference to N^361-369^, bound to HLA*A11:01 weakly (Figure 4B). Meanwhile, IFN-γ production was evaluated for the reactivity of TCR 1- and TCR 4-CD8^+^ T cells, after co-culturing TCR-transduced cells with HLA-A*11:01^+^ COS-7 cells loaded with each mutant peptide. We observed that N^361-369^-specific TCR-T cells responded differently to these mutant peptides. The T366I, F363L or T362I mutations of N^361-369^ could still induce high levels of IFN-γ production, but not other point mutations (Figure 4B). Interestingly, we found that N^T366I^ preferably led to an activation of the TCR 1-T cells similar to that of the wild type N^361-369^, and the TCR 4-T cells were mainly activated by N^T362I^ or N^F363L^ (Figure 4B). These findings indicated that N^361-369^-specific TCRs derived from the COVID-19 convalescent patients had cross-reactivity to certain N^361-369^ variants. Altogether, the N^361-369^ is shown to be a T cell epitope with strong immunogenicity that can be recognized by a diversity of TCRs with high affinity. Such diverse TCRs may play critical protective roles in SASR-CoV-2 infection and decrease the mutation driven immune escape.

### Confirmation of the Endogenous Presentation of the N^361-369^ Epitope

Furthermore, we studied whether the bioinformatically predicted N^361-369^ epitope could be naturally processed and presented from SARS-CoV-2 N protein. First, as dendritic cells (DCs) can ingest virus-infected cells or cellular fragments and cross-present antigens to activate CD8^+^ T cells, we tested whether the N^361-369^ specific-TCR-T cells could be activated by the N protein loaded mature DCs that were induced from the HLA-A*11:01^+^ healthy donors’ monocytes. Substantial levels of T cell activation were detected in TCR 1- and TCR 4-CD8^+^ T cells, indicated by elevated CD137 expression, comparing to the S protein loaded groups (Figure 5A). Also, we found significantly higher levels of IFN-γ secretion in the N protein loaded groups, relative to the S protein controls (Figure 5B).

**Figure 5.**
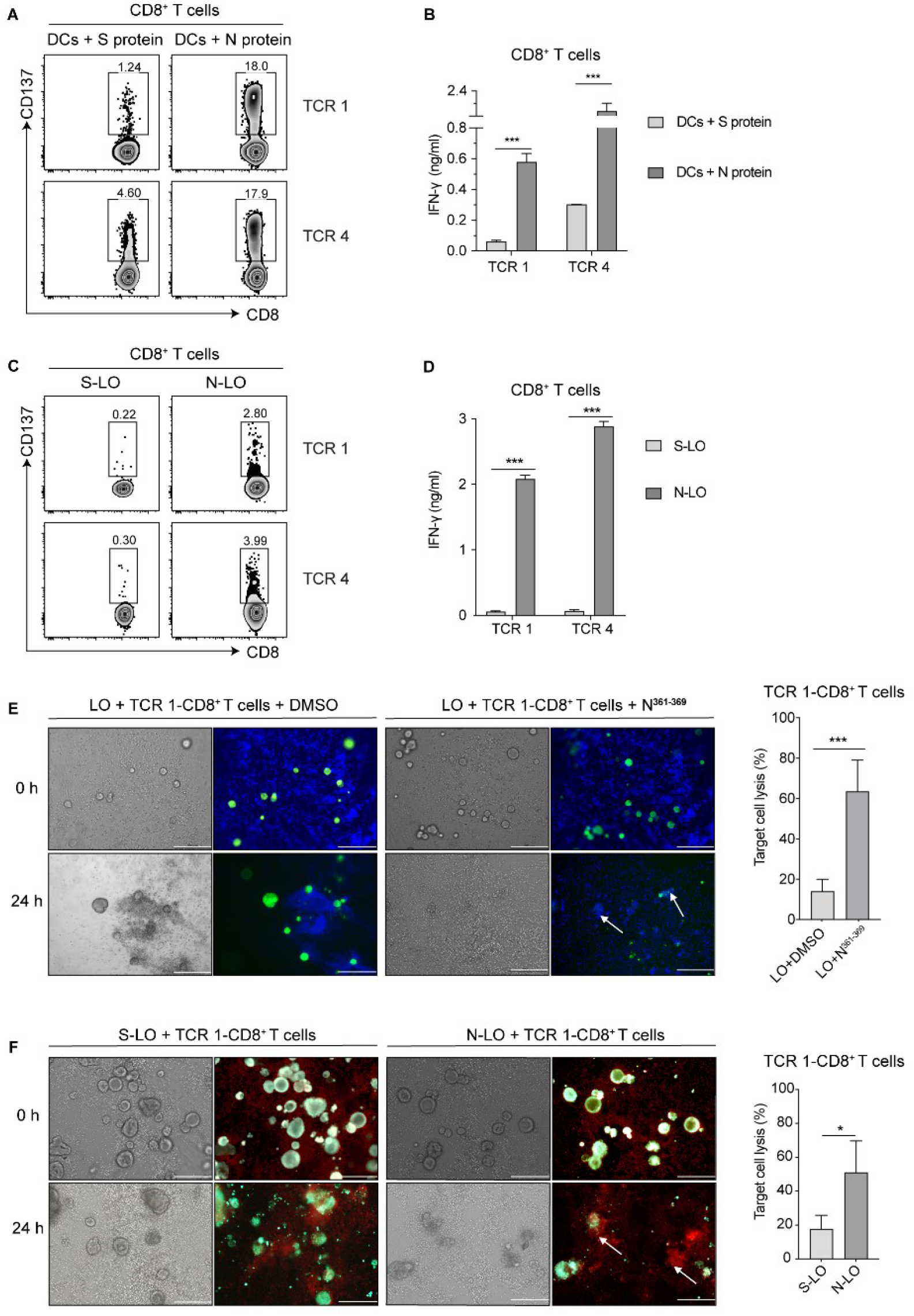
The Reactivity of TCR-Transduced CD8^+^ T Cells to the Endogenously Presented N361-369 Epitope on APCs. The reactivity assay of TCR 1- and TCR 4-transduced CD8^+^ T cells following co-culturing with (A and B) HLA-A*11:01^+^ DCs loaded with N or (C and D) HLA-A*11:01^+^ lung organoids endogenously expressing N protein. (A and C) CD137 expression was evaluated by flow cytometry. (B and D) IFN-γ secretion was determind by ELISA. (E) Cytotoxicity assay of TCR 1-CD8^+^ T cells against HLA-A*11:01^+^ lung organoids exogenously pulsed with N^361-369^ peptide (left). DMSO was used as negative control. T cells were stained with blue dye and lung organoids were stained in green. Target cell lysis was quantified using Calcein-AM release assay (right). (F) Cytotoxicity assay of TCR 1-CD8^+^ T cells against HLA-A*11:01^+^ lung organoids endogenously expressing the N protein (left). Organoids expressing the S protein was applied as negative control. T cells were stained in red and lung organoids were stained in green.Target cell lysis was quantified by LDH release assay (right). Correspoding microphotographs of brightfield displayed all cells in view. White arrows show examples of organoids that are efficiently lysed. Bar: 200 μm. Data were presented as the mean + SD, n = 3, *p < 0.05, ***p < 0.001. Respresentative data of two independent experiments were shown.

Next, we verified whether N^361-369^ could be endogenously presented by lung epithelial cells that were infected by SASR-CoV-2. To this end, we established a series of lung tissue organoids, and HLA-A*11:01^+^ lung organoids, referred as LO, were selected for the subsequent experiments (Figures S5A-5C). The expression of N protein in lung organoid (N-LO) was achieved by ectopically expressing the N protein via the lentiviral EGFP system, with the transfection efficiency confirmed by fluorescent detection (Figure S5D). The lung organoids expressing S protein (S-LO) were used as control. After N^361-369^-specific TCR-T cells were co-cultured with N-LO or S-LO for 24 hours, flow cytometric analysis demonstrated that only N-LO could induce an activation of these TCR-CD8^+^ T cells, indicated by enhanced CD137 expression and significantly elevated IFN-γ production (Figures 5C and 5D). These results demonstrated that the N^361-369^ epitope could be naturally processed from the full length N protein, and presented on the surface of APCs by the intracellular HLA machinery, leading to the activation of N^361-369^ specific CD8^+^ T cells.

To evaluate the N^361-369^ activated CD8^+^ T cells could kill lung organoids, we co-cultured HLA-A*11:01^+^ LO loaded with the N^361-369^ peptide with TCR 1-CD8^+^ T cells. After 24 h, we observed apparent organoid lysis, together with the recruitment of TCR 1-T cells, only with the presence of N^361-369^ (Figure 5E). We found a significantly higher level of target cell lysis in the N^361-369^ loaded group, when these effects were quantified by Calcein-AM release assay (Figure 5E). To determine whether the cytotoxic CD8^+^ T cells could eliminate infected lung organoids, we co-cultured HLA-A*11:01^+^ N-LO or S-LO with TCR 1-CD8^+^ T cells. We found that lung organoids with endogenous expression of the N protein became more susceptible to the cytotoxicity of specific T cells, comparing to the S-LO group (Figure 5F). Such difference of the cytotoxic effect was statistically significant, analysed by LDH release assay (Figure 5F). Taken together, we identified a dominant T cell epitope, N^361-369^, that could activate specific CD8^+^ T cells, and the cytotoxicity of these T cells would result in the lysis of virus infected cells, executively managing the virus clearance in COVID-19 patients.

## DISCUSSION

Currently, limited information is available for the mechanisms of the cellular immunity against SARS-CoV-2. Better understanding of the T cell responses and their protective effects are essential for the vaccine development for COVID-19. Here, we have identified immunodominant epitopes from SARS-CoV-2 S and N proteins that can induce specific CD8^+^ T cell responses, in the majority of COVID-19 convalescent patients with HLA-A*02:01, HLA-A*24:02 or HLA-A*11:01. Importantly, we have discovered N^361-369^, a dominant epitope from the N protein, and 2 different CD8^+^ TCRs recognizing N^361-369^. These 2 TCRs displayed high functional avidity for the wild-type and the mutant variants of N^361-369^, which provided complementary protections for each other against SASR-CoV-2. We are the first to present evidence that N^361-369^ can be endogenously presented by HLA-A*11:01 on APCs, which, in turn, can activate specific CD8^+^ T cells that may contribute to effective virus clearance in COVID-19 convalescent patients.

Both the S and N proteins of SARS-CoV-2 have been reported to effectively eliciting CD8^+^ T cell responses (Grifoni et al., 2020b). Indeed, when stimulated by synthesized peptides corresponding to SARS-CoV-2 S and N, robust CD8^+^ T cell responses could be detected in nearly 75% of COVID-19 convalescent PBMCs (Figure 1; Table S5). Strikingly, we found that 3 out of 4 dominant peptides were derived from the N protein, and accounted for approximately 50% of tested CD8^+^ T cell reactivity. These data suggested that the N protein might be a dominant target for CD8^+^ T cell recognition. Similar observations have been made (Grifoni et al., 2020b; Weiskopf et al., 2020). Of note, T cell responses to the 4 dominant CD8^+^ T cell epitopes identified in our study were not observed in uninfected individuals (Figure S4), while cross-reactivities with other common coronaviruses in unexposed healthy donors had been reported. This difference might due to the refined sample size of healthy donors with corresponding HLA restrictions in our study. Contrast to the rather limited HLA-A phenotype information associated with available T cell epitopes from COVID-19 samples, we provided 15 SARS-CoV-2 epitopes for the 3 most prominent HLA class I A alleles in the Asian population (Table 1), offering more accurate information for vaccine designs. Of these dominant epitopes, the HLA-A*11:01-restricted N^361-369^ (KTFPPTEPK) was found in 5/8 COVID-19 PBMCs (Figure 1C; Table 1). Independently, this epitope has also been identified in multiple individuals in the British COVID-19 convalescent patients (Peng et al., 2020). These findings strongly suggested that N^361-369^ was highly immunogenic and capable to induce robust CD8^+^ T cell responses to SARS-CoV-2.

Presently, reports for SARS-CoV-2 inducing cellular immune responses were largely focused on the T cell-antigen reactivity. Detailed mechanisms regarding the cellular immunity in COVID-19 convalescent patients were needed for advancing the vaccine development and evaluation. In our study, we found 2 N^361-369^-specific TCRs with distinct functions. When ectopically transduced, they could elicit CD8^+^ T cell cytotoxicity upon target cells (Figure 3D), and mediate the activation of T helper type 1 CD4^+^ T cells to secret cytokines in a CD8-independent manner (Figure 3E). It has been reported that such independency could only be achieved when the TCR exhibited high-affinity binding to the cognate pMHC (Campillo-Davo et al., 2020; Lyons et al., 2006; Tan et al., 2017). In the affinity test of the N^361-369^-specific TCRs in the transduced CD8^+^ T or CD4^+^ T cells, we found that their EC50 concentrations ranged from 47.17 nM to 380.9 nM (Figure 4A). These values are remarkably smaller than most of the reported thresholds of affinity for maximal T-cell activity to peptide epitopes, which were around 5-10 μM (Oren et al., 2014; Schmid et al., 2010; Tan et al., 2015; Zhong et al., 2013). Hence, our data evidenced that the N^361-369^-specific TCRs had high affinity and functional avidity, both of which were key features to induce sufficient levels of T cell activation for antivirus responses. As a newly emerged single stranded RNA virus belonging to the highly contagious coronavirus subfamily, SARS-CoV-2 has a tendency to accumulate mutations over time (Duffy, 2018; Villa et al., 2020). Here, another innovative piece of our finding was that the 2 distinct N^361-369^-specific TCRs could recognize multiple N^361-369^ mutations in complementary (Figure 4B). Thus, our findings have revealed the high functional avidity of the 2 TCRs for N^361-369^, with compatible characteristics for clinical mutants of this epitope, providing practical tools to interpret potential mechanisms of CD8^+^ T cell responses against SARS-CoV-2, which are highly relevant in vaccine design and evaluation.

Viral antigens can be phagocytosed and cross-presented by DCs to prime CD8^+^ T cells, or directly processed and presented by infected cells to activate cellular immune responses. Having proven the strong immunogenicity of N^361-369^ *in vitro*, it is of physiological importance to confirm that the N protein can be processed and presented naturally. We proved that N^361-369^ could be presented by DC cells and elicited robust activation of specific CD8^+^ T cells (Figures 5A and 5B). Similarly, marked CD8^+^ T cell cytotoxicity were observed with lung organoids either exogenously loaded or endogenously expressing the N^361-369^ epitope (Figures 5C-5F). The significance of CD8^+^ T cell response for viral clearance has been demonstrated in SARS-CoV infection (Channappanavar et al., 2014; Zhao et al., 2010). Therefore, our findings from DCs and the lung organoid model have, for the first time, revealed potential mechanisms of cellular immunity to eliminate SARS-CoV-2 infected lung cells.

At present, majority of COVID-19 vaccine candidates under development focus on SARS-CoV-2 S protein as the antigen, due to its location and functional significance in virus entry (Chandrashekar et al., 2020; Corbett et al., 2020b; Keech et al., 2020; Mercado et al., 2020a; Yang et al., 2020; Yu et al., 2020; Zhu et al., 2020). Hereby, we have presented evidences of 4 dominant SARS-CoV-2 epitopes, most of which were derived from the N protein, highlighting the potential benefit of SARS-CoV-2 N in vaccine optimization. It should be mentioned that the sequence of the N^361-369^ epitope is identical to a fragment of the SARS-CoV N protein, as this region of the two coronaviruses share more than 90% homology (Grifoni et al., 2020a). In fact, of the total 15 SARS-CoV-2 T cell epitopes presented in our study, 8 were independently confirmed with amino acid sequences identical to SARS-CoV (Tables 1 and S1), indicating probable cross-protective effects of vaccines developed according to these CD8^+^ T cell epitopes.

A larger sample size with subgrouping of COVID-19 patients based on the intensities of their diseases may better represent epitope diversities and be advantageous to understand the correlations between various T cell immune responses and the disease severities. In addition to bioinformatics predicted peptide pools, the application of peptide libraries covering the entire SARS-CoV-2 protein sequences can add detailed resolution of the SARS-CoV-2-specific T cell responses and is currently under investigation. The magnitude and breadth of the observed cytotoxic effect of CD8^+^ T cells may be subjected for debate, and potential undesired pathological damage resulted from the T-cell immunity should be addressed in future studies.

In conclusion, this study has demonstrated potent and broad SARS-CoV-2-specific CD8^+^ T cell responses in the majority of COVID-19 convalescent patients. These confirmed immunodominant epitopes and the identified TCRs provide critical knowledge to understand the cellular immune response to SARS-CoV-2 and the mechanisms for eliminating infected cells, and revealed valuable candidates for vaccine designs.

## ACKNOWLEDGMENTS

We acknowledge the clinical laboratories of Yongchuan Hospital of Chongqing Medical University and the Third Affiliated Hospital of Chongqing Medical University for providing blood samples. We also thank all healthy individuals participated in this study. We appreciate the Department of Immunology of Toyama University for kindly providing technical advices. This study was supported by the Emergency Project from Chongqing Medical University and Chongqing Medical University fund (X4457) with the donation from Mr Yuling Feng.

## AUTHOR CONTRIBUTIONS

A.J., and D.P. conceived and designed the study. Q.C., F.G., Y.W., T.L., S.L., J.H., and J.W. were responsible for collecting samples. C.H., M.S., X.H., Q.C., L.L., S.C., J.Z., J.Z., D.C., and Q.W. performed the experiments. C.H., L.L., and X.H. performed the single cell TCR cloning. M.S., and J.Z. established the lung organoids model. M.S., X.H., C.H., W.W., and K.T. performed data analysis and incorporation. C.H., A.J., and W.W. wrote the manuscript.

## MATERIALS AND METHODS

### Ethics Statement

This project “Screening of SARS-CoV-2-specific T cell epitopes” was approved by the ethics committee of Chongqing Medical University. Informed consents were obtained from all participants.

### Cell Culture

Lenti-X 293T cells was purchased from Takara Biomedical Technology, COS-7 and K-562 cells were purchased from American Type Culture Collection (ATCC). Lenti-X 293T and COS-7 cells were cultured in Dulbecco’s modified Eagle’s medium (DMEM, Thermo Fisher Scientific, USA) supplemented with 10% Fetal Bovine Serum (FBS, Thermo Fisher Scientific, USA), 100 IU/ml penicillin and 100 μg/ml streptomycin (Gibco, USA). K-562 cells were cultured in RPMI-1640 (Thermo Fisher Scientific, USA) with 10% FBS, 100 IU/ml penicillin and 100 μg/ml streptomycin. All cells were maintained at 37°C in an incubator with 5% CO_2_.

### PBMCs Isolation

Blood samples of COVID-19 convalescent patients (Table S3) were obtained from Yongchuan Hospital of Chongqing Medical University and the Third Affiliated Hospital of Chongqing Medical University of Chongqing Medical University. Additionally, blood samples of healthy individuals were obtained from Chongqing Medical University (Table S4). Samples were collected and processed as described before (Weiskopf et al., 2013). All blood was collected in ethylene diamine tetra acetic acid (EDTA) tubes and stored at room temperature prior to processing for PBMCs isolation and plasma collection. The whole blood was centrifuged for 15 min at 1850 rpm to separate the cellular fraction and plasma. The plasma was then carefully separated from the cell pellets and stored at −20℃. The PBMCs were isolated via density gradient centrifugation (Lymphoprep, STEMCELL Technologies, Canada). Isolated PBMC were cryopreserved and stored in liquid nitrogen until used in the assays.

### Preparation and Maturation of DCs

Monocytes were enriched from the PBMCs of healthy individuals by using the EasySep CD14 positive selection kit according to the manufacturer’s instructions (STEMCELL Technologies, Canada). Monocytes were cultured in the DC medium for 6 days, which was made with RPMI-1640 medium supplemented with 10% FBS, 100 IU/ml penicillin, 100 μg/ml streptomycin, 2 mM L-glutamine (Gibco, USA), 800 IU/ml GM-CSF (PeproTech, USA) and 800 IU/ml IL-4 (PeproTech, USA). On day 3, fresh DC media were added to the cultures. On day 6, cells were cryopreserved and stored in liquid nitrogen until used in the assays.

### Lung Organoid Culture

All lung tissues used in this study were obtained after surgical resections from volunteers, and processed for organoid culture within 4 hours. Normal lung tissue organoids were established as reported (Sachs et al., 2019). Briefly, a piece of lung tissue was minced, washed with 10 ml Advanced DMEM/F12 (Thermo Fisher Scientific, USA) containing 1× Glutamax (Thermo Fisher Scientific, USA), 10 mM HEPES (Thermo Fisher Scientific, USA) and antibiotics, termed as AdDF+++, and digested in 10 ml AO medium (Table S8) containing 1 μg/ml collagenase (Sigma-Aldrich, USA) on a shaker at 37°C for 1 hour. The digested lung tissue suspension was sheared using 5 ml plastic Pasteur pipettes before straining over a 100 μm filter. The retained tissue pieces were resuspended in 10 ml AdDF+++ following another shearing step. Strained suspensions were combined and centrifuged, and the pellet was resuspended in 10 ml AdDF+++ and washed once. Lung cell pellets were resuspended in 10 mg/ml cold Cultrex growth factor reduced basement membrane extract (BME), type 2 (Trevigen, USA). Drops of 40 μl BME-cell suspension were added into 24-well plates and solidified at 37°C for 10-20 min. Then 400 μl of AO medium was added to each well and culture in the 37°C incubator with 5% CO_2_. The medium was changed every 4 days and the organoids were passaged every 14-18 days.

### Peptide Library

To define the SARS-CoV-2-specific CD8^+^ T cell epitopes of the S and N proteins derived from SARS-CoV-2 reference sequences (GenBank: MN908947). The NetMHCpan 4.0 algorithm (Jurtz et al., 2017) was used to analyze the binding of 9-mer peptides derived from the full lengths of SARS-CoV-2 S and N proteins, with HLA-A*02:01, HLA-A*24:02 or HLA-A*11:01. For each allele, binding peptides with high strength (Rank Threshold < 0.5) were selected, and a total of 72 predicted epitopes were generated. Additional 6 SARS-CoV epitopes were selected (Ahmed et al., 2020; Grifoni et al., 2020a) (Table S1). Individual peptide’s HLA-A specificity was listed in Table S2. All 78 epitopes were synthesized (GenScript) and dissolved in the corresponding solvents for the subsequent experiments. Detailed information for peptide grouping was summarized in Table S2.

### HLA-A Typing for COVID-19 Convalescent Patients and Healthy Individuals

To screen the HLA-A*11:01 phenotypes of the COVID-19 convalescent patients and health individuals, we used Dynabeads™ mRNA DIRECT™ Purification kit (Thermo Fisher Scientific, USA) to extract RNAs from the PBMC samples. Then, reverse transcription and PCR were performed to amplify HLA-A molecules, with available PCR primer information (Simon et al., 2014). The PCR products were ligated into the T vector (TransGen Biotech, China) and then transformed into the Trans5α competent cells (TransGen Biotech, China). Single bacterial clones of each sample were selected for PCR verification and sequencing. The sequencing results were analyzed in the IMGT/HLA (https://www.ebi.ac.uk/ipd/imgt/hla/) website.

The HLA-A2 and HLA-A24 typing of the COVID-19 convalescent patients and health individuals were analyzed using flow cytometry. For staining, 5×10^5^ PBMCs were resuspended in PBS with 2% FBS (FACS buffer), and stained with the HLA-A2-PE (MBL, Clone BB7.2, K0186-5, Japan) or HLA-A24-PE (MBL, Clone 17A10, K0208-5, Japan) for 30 min at room temperature in the dark. Samples were washed twice with FACS buffer following staining. After the final wash, cells were resuspended in 200 μl FACS buffer. All samples were acquired on a FACSCelesta cytometer (BD Biosciences, USA). The data were analyzed using FlowJo software (TreeStar Inc, USA).

### *In vitro* Stimulation of IFN-γ-producing CD8^+^ T Cells with Selected Peptides

PBMCs were plated in 24-well plates at 2×10^6^ cells per well with 100 IU/ml IL-2 (PeproTech, USA) in the complete medium, which was RPMI-1640 supplemented with 10% FBS, 2 mM L-glutamine, 25 mM HEPES and 10 μg/ml gentamicin. The cells were subsequently stimulated with indicated pools of predicted SARS-CoV-2 peptides, at 5 μM for each peptide. Cells were cultured for 10 days to expand specific T cell populations to corresponding stimulations. Half of the media were refreshed every 3 days, or as required. At day 10, cells were harvested and tested for peptide-specific CD8^+^ T cells by the IFN-γ release ELISPOT assay. The remaining cells were cryopreserved and stored in liquid nitrogen until used in assays.

### IFN-γ ELISPOT Assays

IFN-γ ELISPOT assays were performed as reported and with minor modification (Le Bert et al., 2020b). Briefly, ELISPOT plates (Millipore, USA) were coated with human IFN-γ antibody (Clone 1-D1K, 2 μg/ml, Mabtech, Sweden) overnight at 4°C. T cells were rested in cytokine-free media overnight. Then 2×10^4^ T cells were seeded per well in ELISPOT plates and stimulated for 24 h with pools of predicted SARS-CoV-2 peptides (5 μM each). Stimulation with an equimolar amount of DMSO was performed as the negative control, and Phorbol 12-myristate 13-acetate (PMA)/Ionomycin (P/I) (Sigma-Aldrich, USA) was used as the positive control. Subsequently, the plates were developed with human biotinylated IFN-γ detection antibody (Clone 7-B6-1, 1 μg/ml, Mabtech, Sweden), followed by incubation with Streptavidin-AP (1:1000, Mabtech, Sweden) and BCIP/NBT-plus substrate (Mabtech, Sweden). IFN-γ spots were quantified with the AID ELISPOT Reader (AID, Germany). To quantify positive peptide-specific responses, with 2 × mean spots of the unstimulated wells subtracted from the peptide-stimulated wells, and the results expressed as IFN-γ spots/2×10^4^ PBMCs.

### Flow Cytometry Analysis

For staining, 5×10^5^ cells were re-suspended in FACS buffer, and stained with the anti-human antibody cocktail for 30 min at room temperature in the dark. CD3-BV510 (Clone SK7), CD4-PerCP-Cy5.5 (Clone RPA-T4), CD8-FITC (Clone HIT8a) and CD137-BV421 (Clone 4B4-1) were used for the activation induced T cell marker assay. The P64-tetramer-PE used for the identification of antigen-specific CD8^+^ T lymphocytes was generated using the PE QuickSwitch™ Quant HLA-A*11:01 Tetramer Kit (MBL, tb-7305-k1, Japan). The IFN-γ Secretion Assay Detection Kit (MACS, 130-090-762, Germany) was used for labeling IFN-γ secreting T cells, which were sorted into DNase and RNase free 96-well PCR plates (Bio-Rad, USA) at a single cell level by FACSAria III (BD Biosciences, USA).

Cytokine production was assessed by intracellular cytokine staining (ICS). Briefly, after target cells and effector cells were co-cultured in the well of a 96-well plate for 2 h, both Monensin (1:1000, Biolegend, USA) and Brefeldin A (1:1000, Biolegend, USA) were added to the culture. After 6 h stimulation, cells were washed with FACS buffer, and stained for cell surface markers (described above). Cells were then washed with FACS buffer twice following fixation and permeabilization. Next, cells were washed with Perm/Wash buffer (Biolegend, USA) twice and stained for 25 min at room temperature with anti-human antibody against IFN-γ (BV421, Clone 4S.B3). Cells were washed twice with Perm/Wash buffer and resuspended in FACS buffer, then data was acquired via the FACSCelesta cytometer and analyzed using the FlowJo software.

### HLA Binding Assay

The HLA binding assay was performed using the PE QuickSwitch™ Quant HLA-A*11:01 Tetramer kit following the manufacturer’s instructions. Each lyophilized peptide was dissolved in 1 mM recommended solution. In each well of a 96-well plate, the following materials were added sequentially and gently mixed via pipetting: 25 μl QuickSwitch™ Tetramer, 0.5 μl Peptide Exchange Factor and 0.5 μl of 1 mM peptide. Then, the plate was incubated for 4 h at room temperature while protected from light. The tetramer’s exchange rates were measured by the BD FACSCelesta and calculated by QuickSwitch™ Calculator (https://www.mblintl.com/quickswitch-peptide-exchange-calculator/).

### T Cell Receptor Sequencing and Analysis

For single T cell PCR, the TCRα and TCRβ chain genes were amplified as previously described (Hamana et al., 2016), with minor modifications. Briefly, the reverse transcription (RT) reaction was performed in a two-step method following the PrimeScript™ II Reverse Transcriptase kit protocol (Takara, Japan). For the first step, single-cell sorted plates were thawed on ice and added with 5 μl mix containing dNTP mixture and the primers for TCRα and TCRβ, then incubated at 65℃ for 5 min and put on ice immediately. For the second step, the above plates were added with the RNase inhibitor and PrimeScript II Reverse Transcriptase (Takara, Japan) to a total volume of 10 μl. The RT program was 45℃ for 45 min and 70℃ for 15 min. The RT products were amplified by nested PCR following the PrimeSTAR® HS DNA Polymerase kit protocol (Takara, Japan), with primers for TCRα and TCRβ. Here, the 5×PrimeSTAR^®^ Buffer (Mg^2+^ plus) in the kit was replaced with the 2 × PrimeSTAR^®^ GC Buffer (Mg^2+^ plus) (Takara, Japan). The first PCR program was as follows: 98℃ for 1 min, 98℃ for 10 sec for 30 cycles, 52℃ for 5 sec and 72℃ for 40 sec. The resultant PCR mixtures were diluted 50-fold with water, and 2 μl of the diluted PCR mixtures were added to 18 μl of the nested PCR mixture as the template DNA. The nested PCR program was as follows: 98℃ for 1 min, 98℃ for 10 sec for 35 cycles, 52℃ for 5 sec and 72℃ for 30 sec. The PCR products were then analyzed with the α-nest or β-nest primers by direct sequencing. The TCR repertoire was analyzed with the IMGT/V-Quest tool (http://www.imgt.org/) (Brochet et al., 2008).

### Enrichment of CD3^+^, CD4^+^ and CD8^+^ T Cells

CD3^+^, CD4^+^ or CD8^+^ T cells were selected using the EasySep human CD3 positive selection kit II, EasySep human CD4^+^ T cell enrichment kit and EasySep human CD8^+^ T cell enrichment kit (STEMCELL Technologies, Canada), respectively, according to the manufacturer’s instructions.

### Construction of Lentivirus Vectors and Transduction of CD3^+^ T Cells

For the construction of TCR lentivirus vectors, TCRα or TCRβ chains were amplificated from the corresponding nested PCR mixtures and cloned into the lentivirus vector pWPXL (Addgene Plasmid #12257). The constant regions were replaced by mouse counterparts with improved TCR pairing and TCR/CD3 stability that were convenient for the detection of TCR-T cells (Cohen et al., 2006; Jin et al., 2018). The coding sequence of the S or N gene of SARS-CoV-2 and the human *HLA-A*11:01* gene were synthesized, and cloned into the pWPXL vector. Lentiviruses were generated by co-transfecting Lenti-X 293T cells with the lenti-vector and the packaging plasmids psPAX2 (Addgene Plasmid #12260) and pMD2.G (Addgene Plasmid #12259) at a ratio of 5:2:1, using the Xfect Transfection Reagent (Takara, Japan). The lentiviral supernatants were harvested in 48 h. Transduction of CD3^+^ T cells was conducted as delineated before (Jin et al., 2018), with minor modifications. CD3^+^ T cells were stimulated in T cell media with 1 μg/ml anti-CD3 antibody (Clone OKT3) and 1 μg/ml anti-CD28 antibody (Clone 15E8) for 2 days. Then, the transduction efficiency was assessed by flow cytometry using anti-mouse TCR-β chain constant region antibody (Clone H57-597).

### TCR Transduced-T Cell Rapid Expansion Protocol

TCR transduced-T cells were rapidly expanded with the method as described before (Jin et al., 2012). Briefly, in indicated experiments, sorted T cells were expanded using excess irradiated (50Gy) allogeneic PBMC feeder cells (a pool from 3-5 different donors) at a ratio of 1 to 100 with 30 ng/ml anti-CD3 antibody (Clone OKT3) and 3000 IU/ml IL-2, in the 50/50 media consisted of a 1:1 mixture of the complete media and the AIM-V media (Gibco, USA). All cells were cultured at 37°C with 5% CO_2_. Media were exchanged 3 days post-stimulation and then every other day. The cells were rapidly expanded in this way for two weeks, before use.

### Antigen Presentation in Human DCs

DCs were incubated in AIM-V for 1 h and then 5 μg/ml recombinant N or S proteins of SARS-CoV-2 were added. After 4 h, the media were replaced to DC media supplemented with DC maturation cytokines including 10 ng/ml Lipopolysaccharide (LPS) (Sigma-Aldrich, USA) and 100 IU/ml IFN-γ (PeproTech, USA). After incubated overnight, DCs were washed and co-cultured with TCR-T cells for 24 h. The supernatants were used for measuring IFN-γ concentrations by ELISA.

### TCR-T Cytotoxicity Assay

The ability of T cells to lyse target cells was measured using a Calcein-AM release assay. Target cells were labelled for 15 min at 37°C with 10 μM Calcein-AM (Dojindo, Japan), and then co-cultured with T cells at indicated ratios for 24 h in the black-walled, 96-well flat-bottom plates (Corning, USA). The controls of these labelled target cells were separated into the spontaneous death group and the maximal killing group, the latter were treated with 0.1% Triton-X (Sigma-Aldrich, USA). After 24 hours, 50 μl/well supernatant was removed, and the relative fluorescence units (RFUs) was measured by Varioskan LUX Multimode Microplate Reader (Thermo Fisher Scientific, USA). The percentage of lysed cells was calculated as follows: specific lysis = 100% × (Test RFUs - Spontaneous death RFUs) / (Maximal killing RFUs - Spontaneous death RFUs).

The ability of T cells to lyse organoids was also measured using LDH-Glo™ Cytotoxicity Assay Kit (Promega, USA) according to the manufacturer’s instructions. The organoids were dissociated to single cells and counted, which was used to infer the number of organoids to allow co-cultured with T cells at a 4:1 effector:target ratio. After co-cultured for 24 h in the 96-well flat-bottom plates, the relative luminescent units (RLUs) of the co-culture and the control groups were measured by the Varioskan LUX Multimode Microplate Reader. The percentage of lysed organoids was calculated as follows: specific lysis = 100% × (Test RLUs - Spontaneous death RLUs) / (Maximal killing RLUs - Spontaneous death RLUs).

### Cytokine Release ELISA

The antibody pairs used for cytokine detection are as follows: IFN-γ capture antibody (Clone MD-1, 2 μg/ml), IFN-γ detection antibody (Clone 4S.B3, 1 μg/ml), IL-2 capture antibody (Clone MQ1-17H12, 4 μg/ml), IL-2 detection antibody (Clone Poly5176, 1 μg/ml), TNF-α capture antibody (Clone MAb1, 1 μg/ml) and TNF-α detection antibody (Clone MAb11, 1μg/ml). The ELISA plate was coated with individual anti-cytokine capture antibodies for 12 h, then washed with phosphate-buffered saline with Tween (PBST) buffer for 3 times, followed by blocking with 3% BSA for at least 1 h at room temperature. The supernatants of T cell and target cell co-cultures were added into the plate at 50 μl per well. After incubation at room temperature for 1 h, the plate was washed 3 times with PBST. Biotinylated capture antibodies (50 μl/well) were added and the plate was incubated for 1 h at room temperature, then washed for 5 times. Streptavidin-ALP (1:1000) was added at 50 μl per well, with, and incubated for 1 h at room temperature in the dark. Then, the plate was washed 6 times and the pNPP solution (Mabtech, Sweden) was added at 50 μl per well. After 20 min incubation at room temperature in the dark, 50 μl stop solution was added per well. The plate was immediately analyzed by the Varioskan LUX Multimode Microplate Reader for the OD405 value.

### Statistical Analysis

Statistical analyses of the data were performed using GraphPad Prism version 8.0 (GraphPad Software, Inc. La Jolla, CA, USA) software. Quantitative data in histograms and line charts were presented as mean ± standard deviation (mean ± SD). Statistical significance was determined using ANOVA for multiple comparisons. Student’s *t*-tests were applied to compare two groups. A value of *P* < 0.05 was considered statistically significant.

---

**Figure S1.**
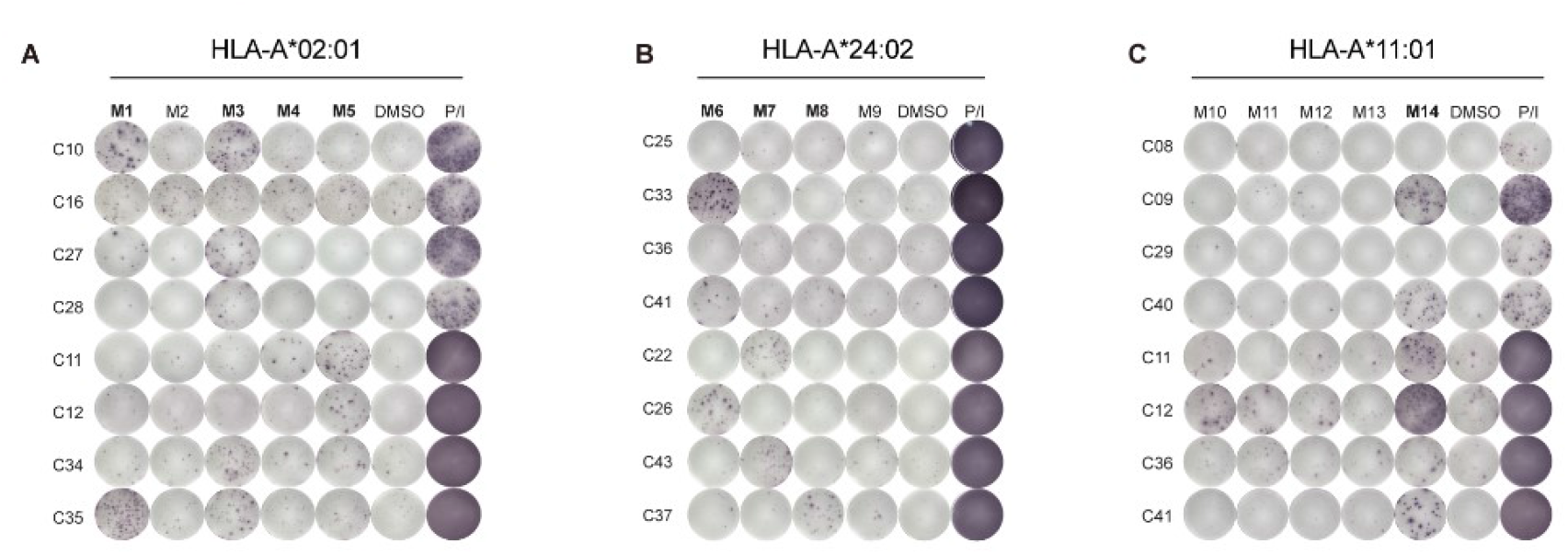
T Cell Responses after Stimulation with Predicted SARS-CoV-2 Epitope Pools. Related to Figure 1. IFN-γ ELISPOT assay results for the PBMCs of COVID-19 convalescent patients with (A) HLA-A2^+^ allele, (B) HLA-A24^+^ allele and (C) HLA-A11:01^+^ allele, stimulated with peptide mixtures. Results are expressed as number of peptide-specific IFN-γ spots/2×10^4^ PBMCs of each patient, substracted those from the corresponding DMSO groups. For peptide grouping: 26 HLA-A*02:01-restricted peptides were pooled into 5 mixtures (M1-M5), 22 HLA-A*24:02-restricted peptides were pooled into 4 mixtures (M6-M9), and 30 HLA-A*11:01-restricted peptides were pooled into 5 mixtures (M10-M14). Each mixture contained 3-6 peptides. Positive peptide mixtures were marked in bold. A stimulation with an equimolar amount of DMSO was performed as the negative control, and phorbol 12-myristate 13-acetate (PMA)/Ionomycin (P/I) was used as the positive control. Data were representative of two independent experiments.

**Figure S2.**
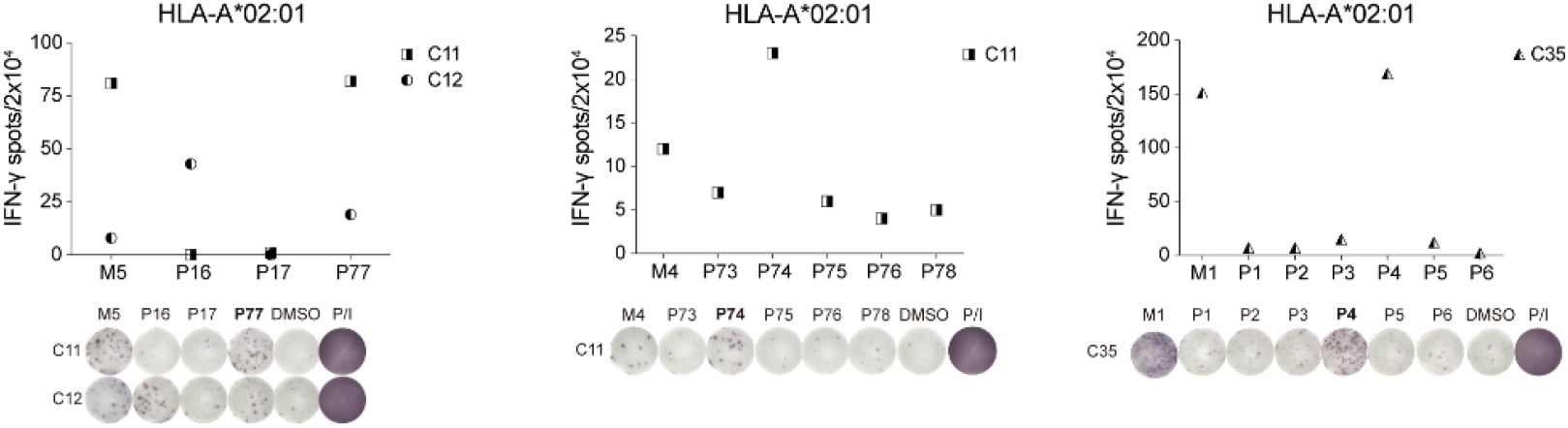
Identification of T Cell Epitopes in HLA-A2+ Patients. Related to Figure 1. Individual image summarized (top) and directly showed (bottom) IFN-γ ELISPOT assay results for PBMCs of COVID-19 convalescent patients with HLA-A2^+^ allele, which were stimulated with a positive peptide mixture and the corresponding individual peptides from the mixture. Results are expressed as number of peptide-specific IFN-γ spots/2×10^4^ PBMCs of each patient, substracted those from the corresponding DMSO groups. DMSO and P/I were used as the negative and positive control respectively. Positive single peoptide was marked in bold. Data were representative of two independent experiments.

**Figure S3.**
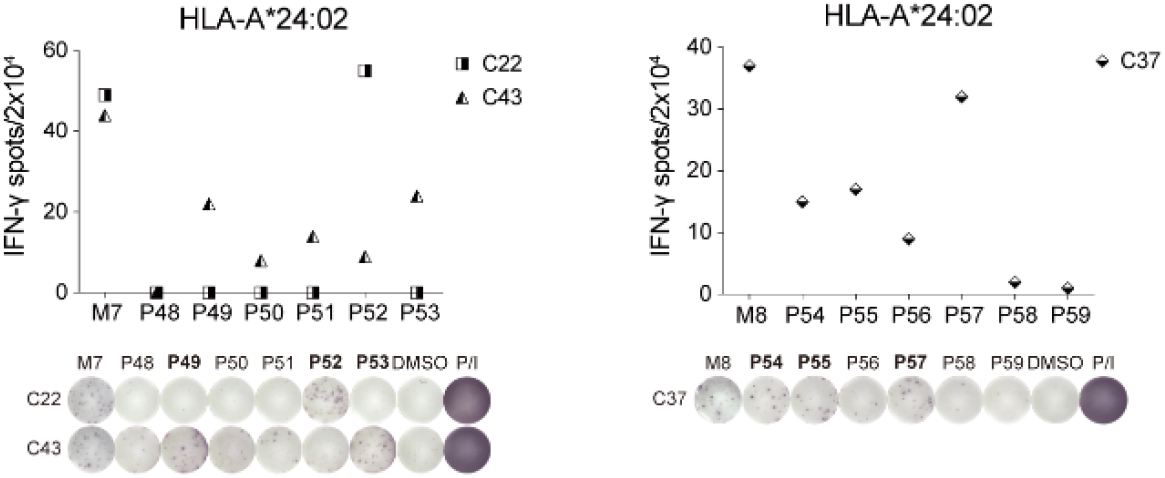
Identification of T Cell Epitopes in HLA-A24+ Patients. Related to Figure 1. Individual image summarized (top) and directly showed (bottom) IFN-γ ELISPOT assay results for PBMCs of COVID-19 convalescent patients with HLA-A24^+^ allele, which were stimulated with the positive peptide mixture and the corresponding 6 individual peptides from the mixture. Results are expressed as number of peptide-specific IFN-γ spots/2×10^4^ PBMCs of each patient, substracted those from the corresponding DMSO groups. DMSO and P/I were used as the negative and positive control respectively. Positive single peoptide was marked in bold. Data were representative of two independent experiments.

**Figure S4.**
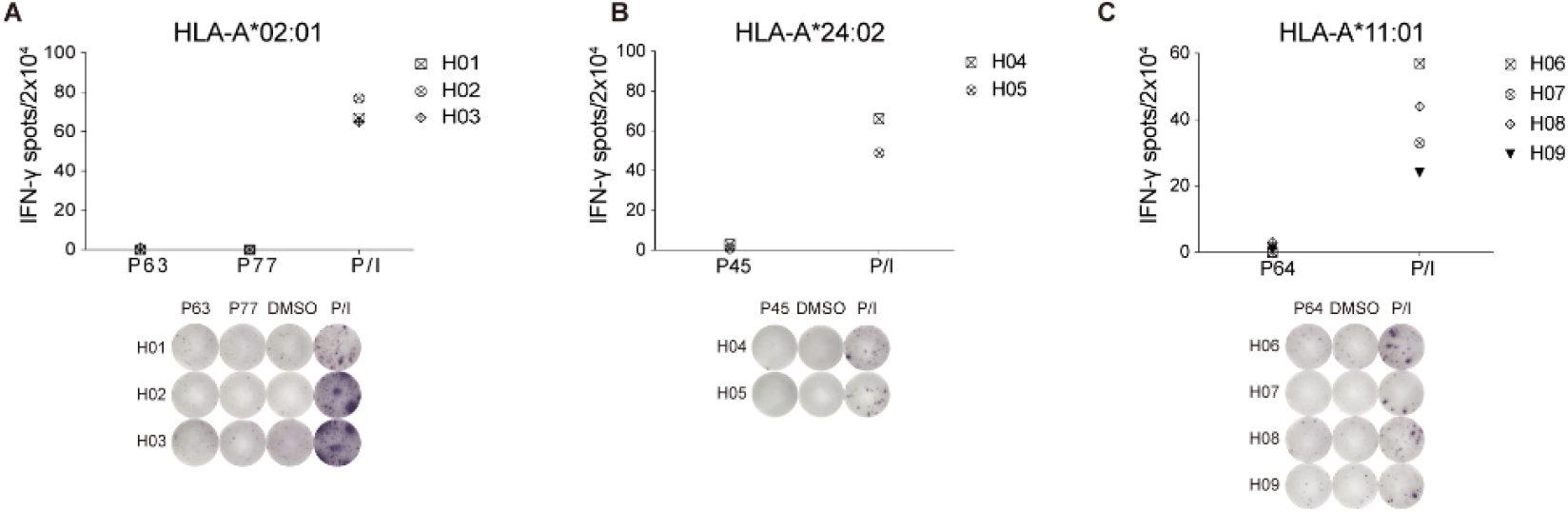
Assessment of SARS-CoV-2 Dominant Epitopes Reactivity in Healthy Individuals. Related to Figure 1. Individual image summarized (top) and directly showed (bottom) IFN-γ ELISPOT assay results for the PBMCs of healthy individuals with (A) HLA-A2^+^ allele, (B) HLA-A24^+^ allele and (C) HLA-A11:01^+^ allele, stimulated with corresponding dominant epitope(s). Results are expressed as number of peptide-specific IFN-γ spots/2×10^4^ PBMCs of each patient, substracted those from the corresponding DMSO group. DMSO and P/I were used as the negative and positive control respectively. Data were representative of two independent experiments.

**Figure S5.**
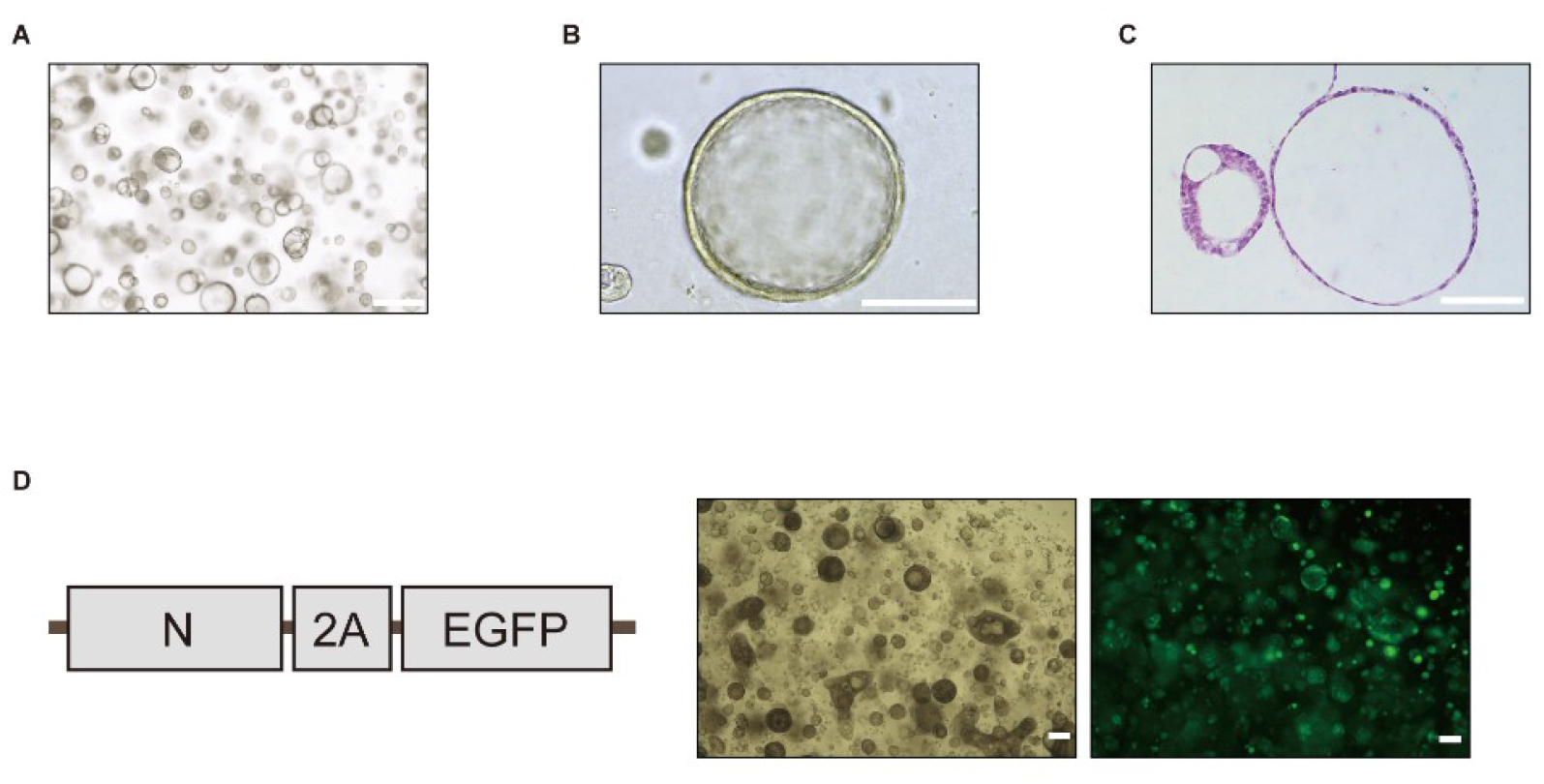
Establishment of the Lung Tissue Organoid Model for the Assessment of SARS-CoV-2 N Antigen Presentation. Related to Figure 5. (A and B) Bright-field microscopy images and (C) H&E staining image of lung tissue organoids cultured for one month. Bar indicates 100 μm. (D) A schematic diagram represents the open reading frame (ORF) of the N protein lentivirus expression vector (left). The coding sequences of N protein, 2A and EGFP were connected in this vector. The transfection efficiency of N protein lentiviruses in lung organiods was examined. EGFP expression of the positively transfected organoid was shown in the fluorescent microphotograph (right).

**Supplementary Table 1.**
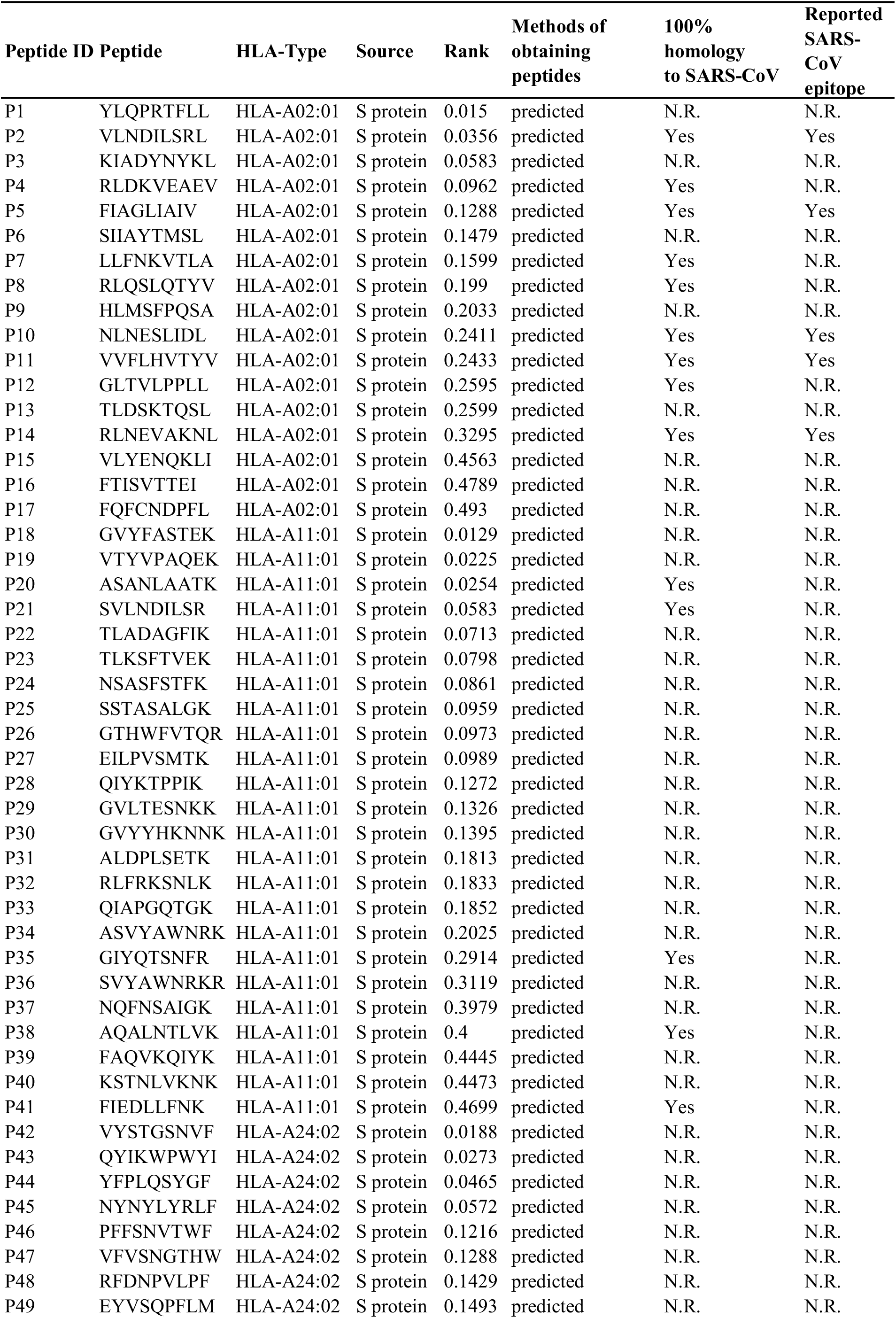
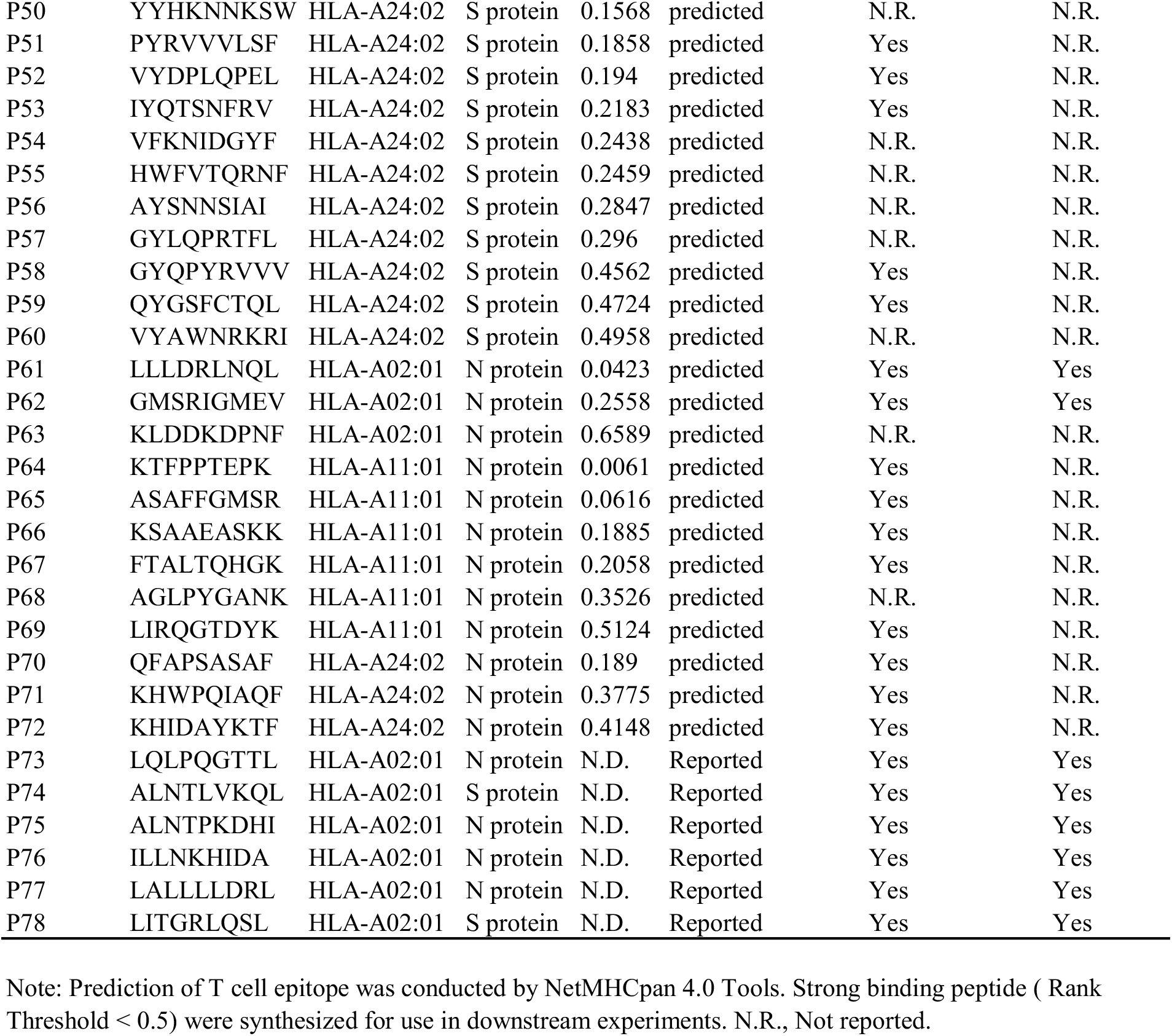
Prediction and Selection of T Cell Epitopes from SARS-CoV-2 S and N Proteins

**Supplementary Table 2.**
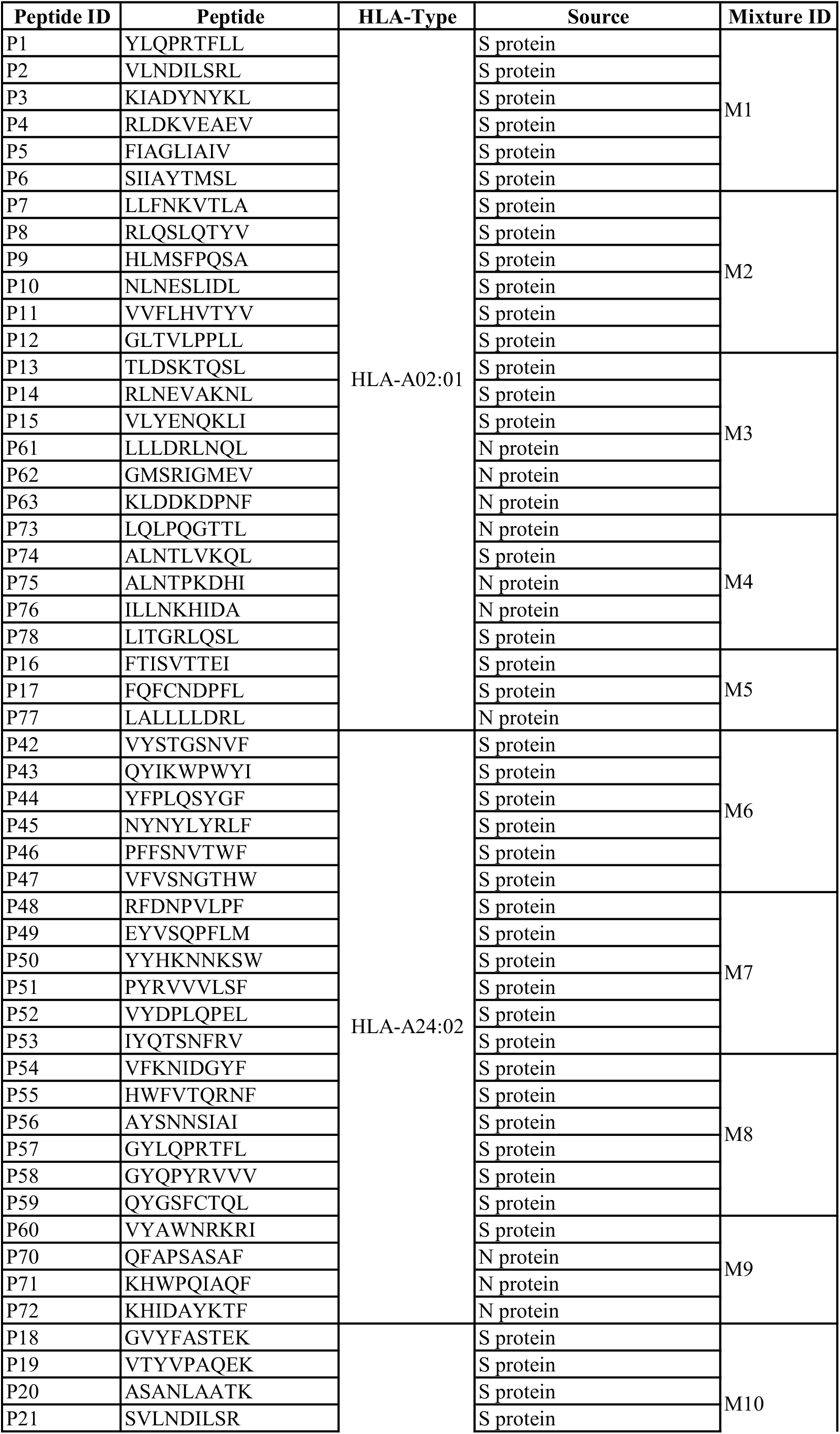
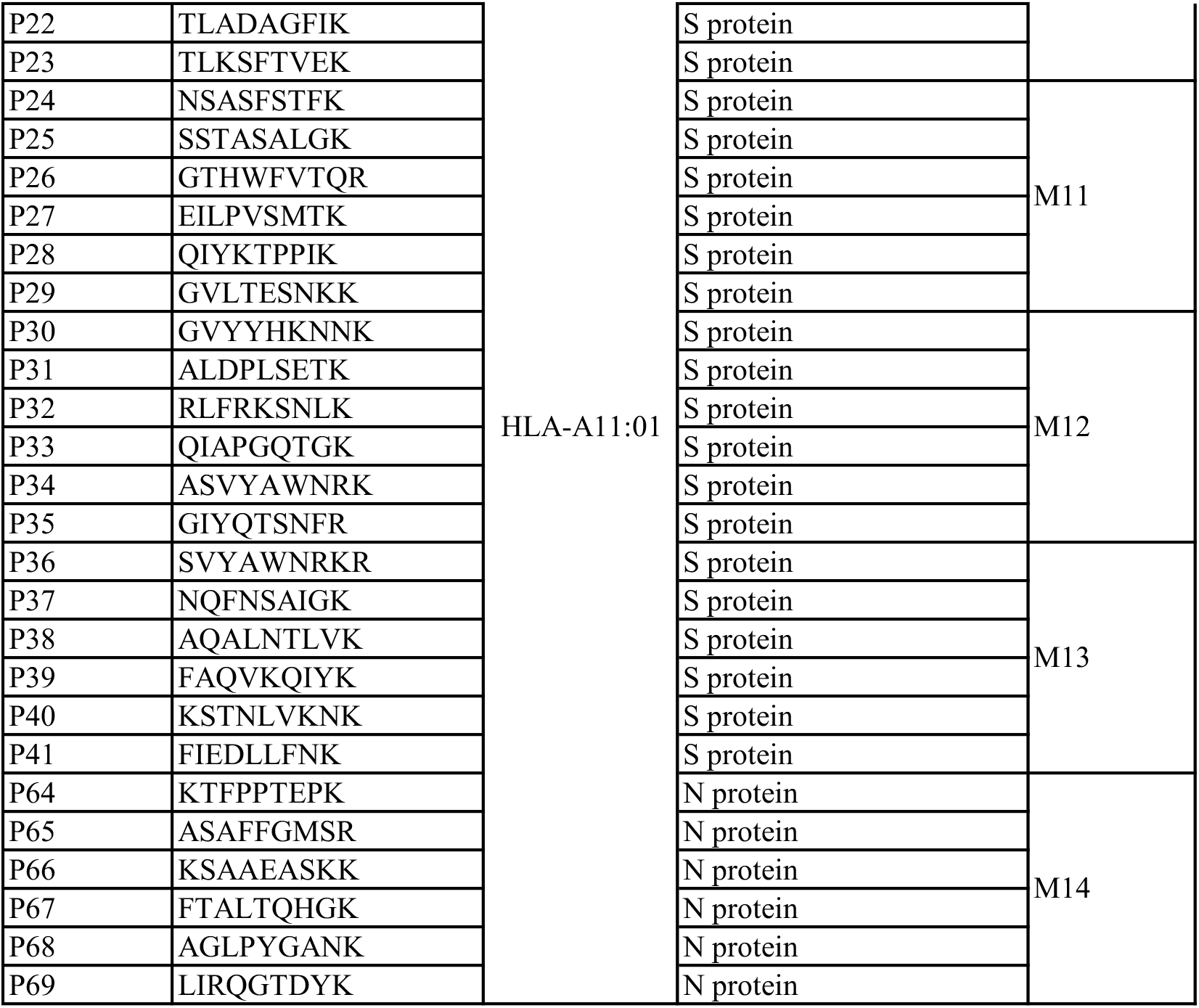
The Grouping Information of Synthetic Peptides

**Supplementary Table 3.** SARS-CoV-2 Convalescent Patient Information

| Patient ID | Age (years) | Sex | Subject type | Citywhere infected | Clinical classification (mild:0, severe:1) | Date of admission | Date of discharge | HLA-A type |  |  | Usage for the epitopes screening |
| --- | --- | --- | --- | --- | --- | --- | --- | --- | --- | --- | --- |
|  |  |  |  |  |  |  |  | A2 | A24 | A11:01 |  |
| C08 | 51 | F | SARS-CoV-2 convalescent patient | Chongqing | 0 | 2020/1/31 | 2020/2/17 | - | - | + | A11:01 epitope screening |
| C09 | 48 | M | SARS-CoV-2 convalescent patient | Chongqing | 1 | 2020/1/30 | 2020/2/18 | - | - | + | A11:01 epitope screening |
| C10 | 32 | M | SARS-CoV-2 convalescent patient | Chongqing | 0 | 2020/1/29 | 2020/2/18 | + | - | N.D. | A02:01 epitope screening |
| C11 | 54 | M | SARS-CoV-2 convalescent patient | Chongqing | 0 | 2020/2/2 | 2020/2/18 | + | - | + | A02:01 epitope screening<br>A11:01 epitope screening |
| C12 | 43 | M | SARS-CoV-2 convalescent patient | Chongqing | 0 | 2020/1/31 | 2020/2/25 | + | - | + | A02:01 epitope screening<br>A11:01 epitope screening |
| C13 | 47 | F | SARS-CoV-2 convalescent patient | Chongqing | 0 | 2020/1/31 | 2020/2/10 | + | - | N.D. | No |
| C14 | 38 | M | SARS-CoV-2 convalescent patient | Chongqing | 0 | 2020/2/10 | 2020/2/19 | + | - | N.D. | No |
| C15 | 33 | F | SARS-CoV-2 convalescent patient | Chongqing | 0 | 2020/2/10 | 2020/3/9 | + | - | N.D. | No |
| C16 | 44 | M | SARS-CoV-2 convalescent patient | Chongqing | 0 | 2020/2/5 | 2020/2/14 | + | - | N.D. | A02:01 epitope screening |
| C17 | 60 | F | SARS-CoV-2 convalescent patient | Chongqing | 0 | 2020/2/11 | 2020/3/5 | N.D. | N.D. | N.D. | No |
| C18 | 48 | F | SARS-CoV-2 convalescent patient | Chongqing | 0 | 2020/2/12 | 2020/3/6 | N.D. | N.D. | N.D. | No |
| C19 | 53 | F | SARS-CoV-2 convalescent patient | Chongqing | 0 | 2020/2/16 | 2020/2/28 | - | - | N.D. | No |
| C20 | 57 | M | SARS-CoV-2 convalescent patient | Chongqing | 0 | 2020/1/29 | 2020/2/14 | + | - | N.D. | No |
| C21 | 44 | M | SARS-CoV-2 convalescent patient | Chongqing | 0 | 2020/2/6 | 2020/2/29 | - | - | N.D. | No |
| C22 | 49 | M | SARS-CoV-2 convalescent patient | Chongqing | 0 | 2020/2/4 | 2020/3/2 | - | + | N.D. | A24:02 epitope screening |
| C23 | 23 | M | SARS-CoV-2 convalescent patient | Chongqing | 0 | 2020/2/6 | 2020/3/9 | - | + | N.D. | No |
| C24 | 44 | F | SARS-CoV-2 convalescent patient | Chongqing | 0 | 2020/2/13 | 2020/3/11 | + | - | N.D. | No |
| C25 | 59 | F | SARS-CoV-2 convalescent patient | Chongqing | 0 | 2020/2/1 | 2020/2/18 | - | + | N.D. | A24:02 epitope screening |
| C26 | 20 | F | SARS-CoV-2 convalescent patient | Chongqing | 0 | 2020/2/15 | 2020/2/25 | + | + | N.D. | A24:02 epitope screening |
| C27 | 37 | M | SARS-CoV-2 convalescent patient | Chongqing | 0 | 2020/2/2 | 2020/3/9 | + | - | N.D. | A02:01 epitope screening |
| C28 | 44 | F | SARS-CoV-2 convalescent patient | Chongqing | 0 | 2020/2/3 | 2020/2/28 | + | - | N.D. | A02:01 epitope screening |
| C29 | 41 | M | SARS-CoV-2 convalescent patient | Chongqing | 0 | 2020/1/31 | 2020/2/16 | - | - | + | A11:01 epitope screening |
| C30 | 66 | M | SARS-CoV-2 convalescent patient | Chongqing | 0 | 2020/2/3 | 2020/2/16 | + | + | N.D. | No |
| C31 | 67 | F | SARS-CoV-2 convalescent patient | Chongqing | 0 | 2020/2/4 | 2020/2/16 | - | + | N.D. | No |
| C32 | 50 | M | SARS-CoV-2 convalescent patient | Chongqing | 1 | 2020/2/1 | 2020/2/29 | - | - | N.D. | No |
| C33 | 50 | F | SARS-CoV-2 convalescent patient | Chongqing | 0 | 2020/1/28 | 2020/2/29 | + | + | N.D. | A24:02 epitope screening |
| C34 | 49 | F | SARS-CoV-2 convalescent patient | Chongqing | 0 | 2020/2/8 | 2020/3/6 | + | - | N.D. | A02:01 epitope screening |
| C35 | 56 | M | SARS-CoV-2 convalescent patient | Chongqing | 0 | 2020/2/1 | 2020/2/9 | + | - | N.D. | A02:01 epitope screening |
| C36 | 55 | M | SARS-CoV-2 convalescent patient | Chongqing | 0 | 2020/2/14 | 2020/2/21 | + | + | + | A11:01 epitope screening<br>A24:02 epitope screening |
| C37 | 45 | F | SARS-CoV-2 convalescent patient | Chongqing | 0 | 2020/2/1 | 2020/2/15 | + | + | N.D. | A24:02 epitope screening |
| C38 | 47 | F | SARS-CoV-2 convalescent patient | Chongqing | 0 | 2020/2/6 | 2020/2/17 | + | + | N.D. | No |
| C39 | 34 | M | SARS-CoV-2 convalescent patient | Chongqing | 0 | 2020/1/28 | 2020/2/7 | - | - | N.D. | No |
| C40 | 47 | M | SARS-CoV-2 convalescent patient | Chongqing | 1 | 2020/1/27 | 2020/2/23 | - | - | + | A11:01 epitope screening |
| C41 | 56 | M | SARS-CoV-2 convalescent patient | Chongqing | 0 | 2020/1/30 | 2020/2/7 | - | + | + | A11:01 epitope screening<br>A24:02 epitope screening |
| C42 | 63 | F | SARS-CoV-2 convalescent patient | Chongqing | 0 | 2020/2/1 | 2020/2/25 | + | - | N.D. | No |
| C43 | 41 | M | SARS-CoV-2 convalescent patient | Chongqing | 0 | 2020/1/30 | 2020/2/18 | - | + | N.D. | A24:02 epitope screening |
| C44 | 37 | M | SARS-CoV-2 convalescent patient | Chongqing | 0 | 2020/1/27 | 2020/2/14 | - | - | N.D. | No |

Note: F, Female. M, Male. N.D., Not detected.

**Supplementary Table 4.** HLA-A Types of Healthy Donors

| Donor ID | HLA-A Type |  |  | Usage for the epitopes screening |
| --- | --- | --- | --- | --- |
|  | A2 | A24 | A11:01 |  |
| H01 | + | - | - | A02:01 epitope screening |
| H02 | + | - | - | A02:01 epitope screening |
| H03 | + | - | - | A02:01 epitope screening |
| H04 | - | + | - | A24:02 epitope screening |
| H05 | - | + | - | A24:02 epitope screening |
| H06 | - | - | + | A11:01 epitope screening |
| H07 | - | - | + | A11:01 epitope screening |
| H08 | - | - | + | A11:01 epitope screening |
| H09 | - | - | + | A11:01 epitope screening |

Note: H, Healthy Donor.

**Supplementary Table 5.** The Reactivity of Identified T Cell Epitopes in COVID-19 Convalescent Patients

| HLA restriction | Number | Patient ID | Reactive peptide |
| --- | --- | --- | --- |
| HLA-A*02:01 | 1 | C10 | P63 |
|  | 2 | C11* | P74, P77 |
|  | 3 | C12* | P16, P77 |
|  | 4 | C16 | N.D. |
|  | 5 | C27 | P63 |
|  | 6 | C28 | P63 |
|  | 7 | C34 | P62 |
|  | 8 | C35 | P4, P61, P63 |
| HLA-A*24:02 | 1 | C22 | P52 |
|  | 2 | C25 | N.D. |
|  | 3 | C26 | P45 |
|  | 4 | C33 | P45 |
|  | 5 | C36* | N.D. |
|  | 6 | C37 | P54, P55, P57 |
|  | 7 | C41* | N.D. |
|  | 8 | C43 | P49, P53 |
| HLA-A*11:01 | 1 | C8 | N.D. |
|  | 2 | C9 | P64 |
|  | 3 | C11* | P64 |
|  | 4 | C12* | P64 |
|  | 5 | C29 | N.D. |
|  | 6 | C36* | N.D. |
|  | 7 | C40 | P64 |
|  | 8 | C41* | P64 |

Note: C, COVID-19 convalescent patient. N.D., Not detected. \*indicates the repeatedly used patient samples.

**Supplementary Table 6.**
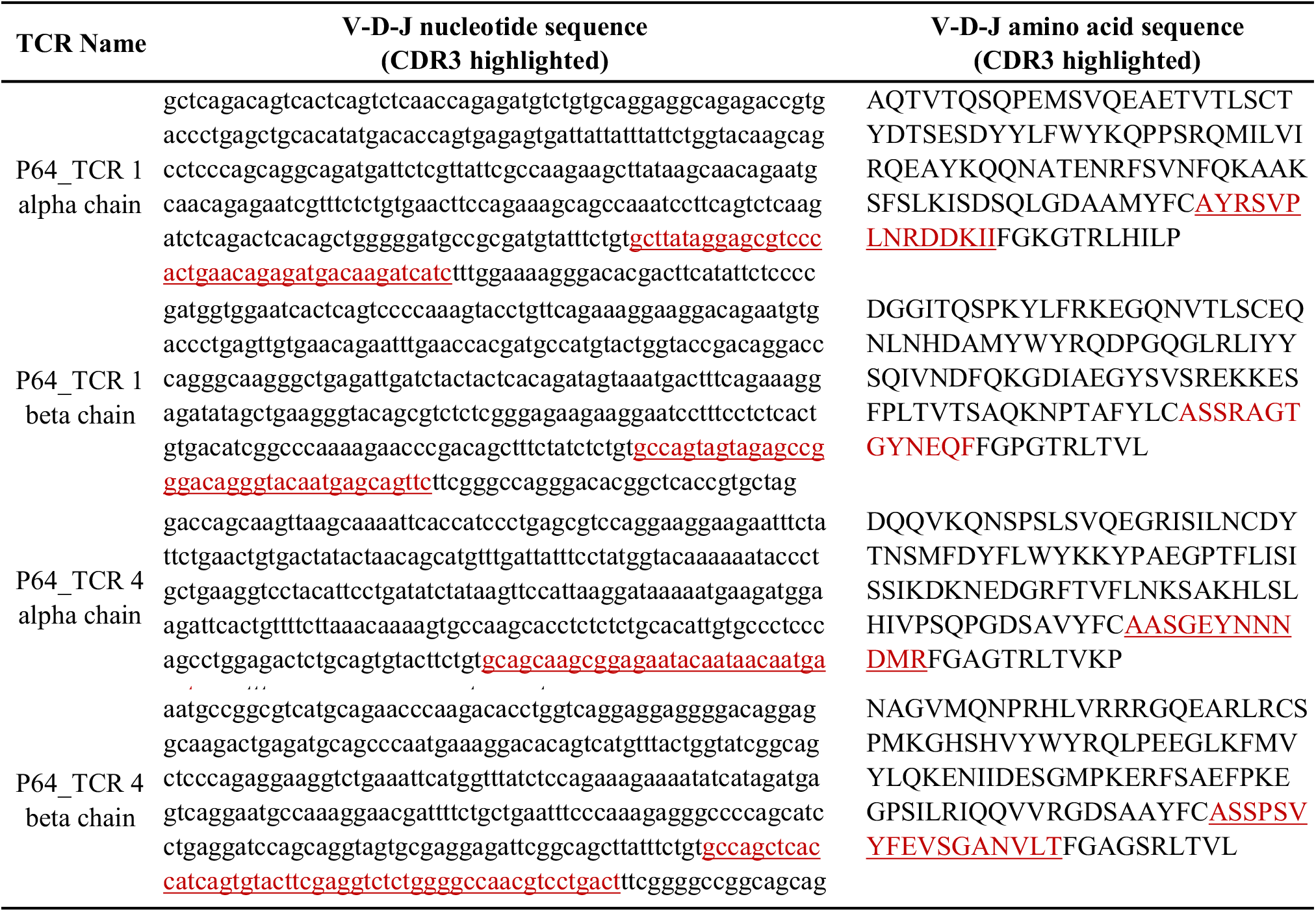
The Sequences of P64 Reactive TCRsementary

**Supplementary Table 7.**
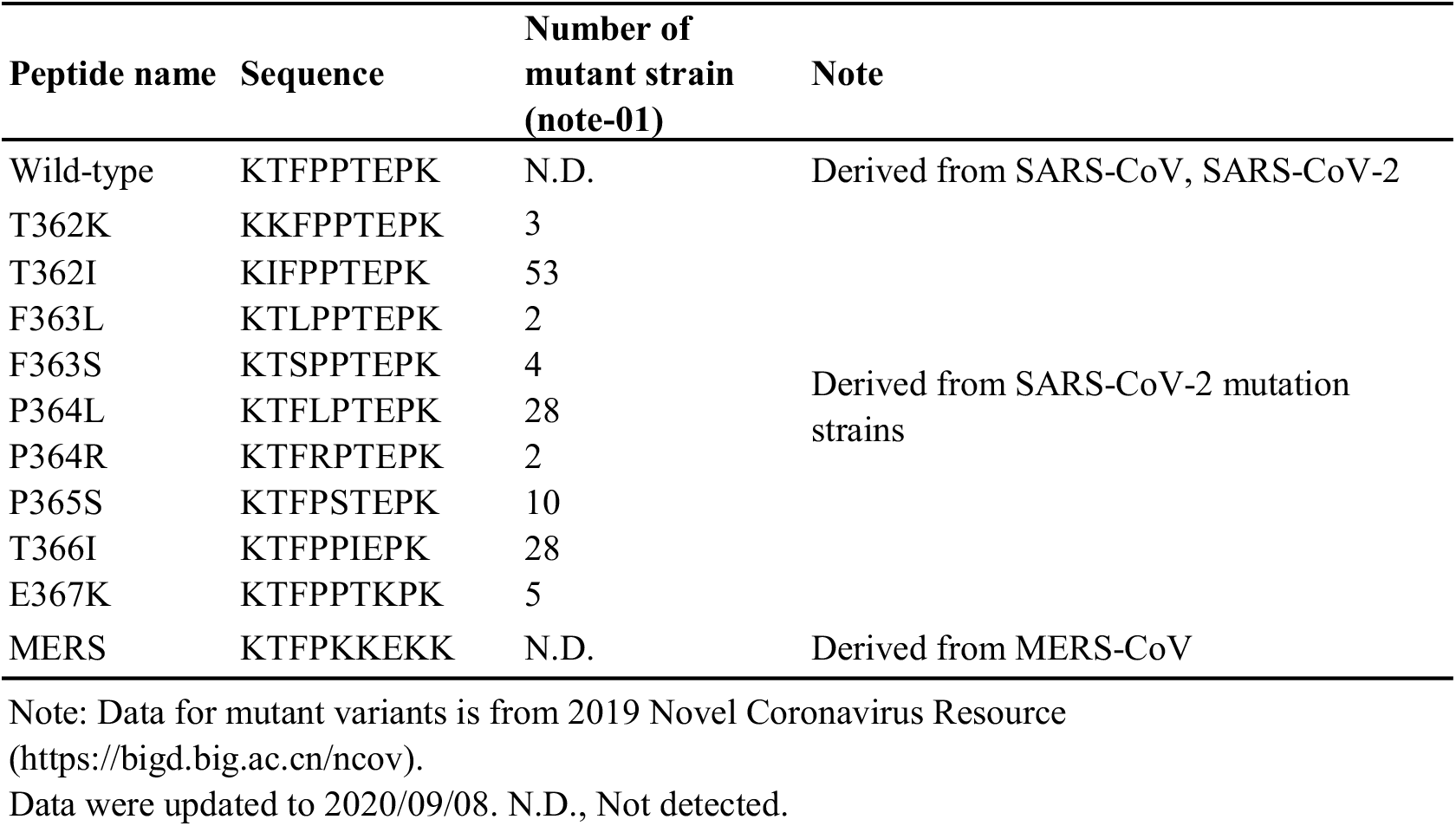
Mutant Variants and Homologous Peptides of N^361-369^

**Supplementary Table 8.**
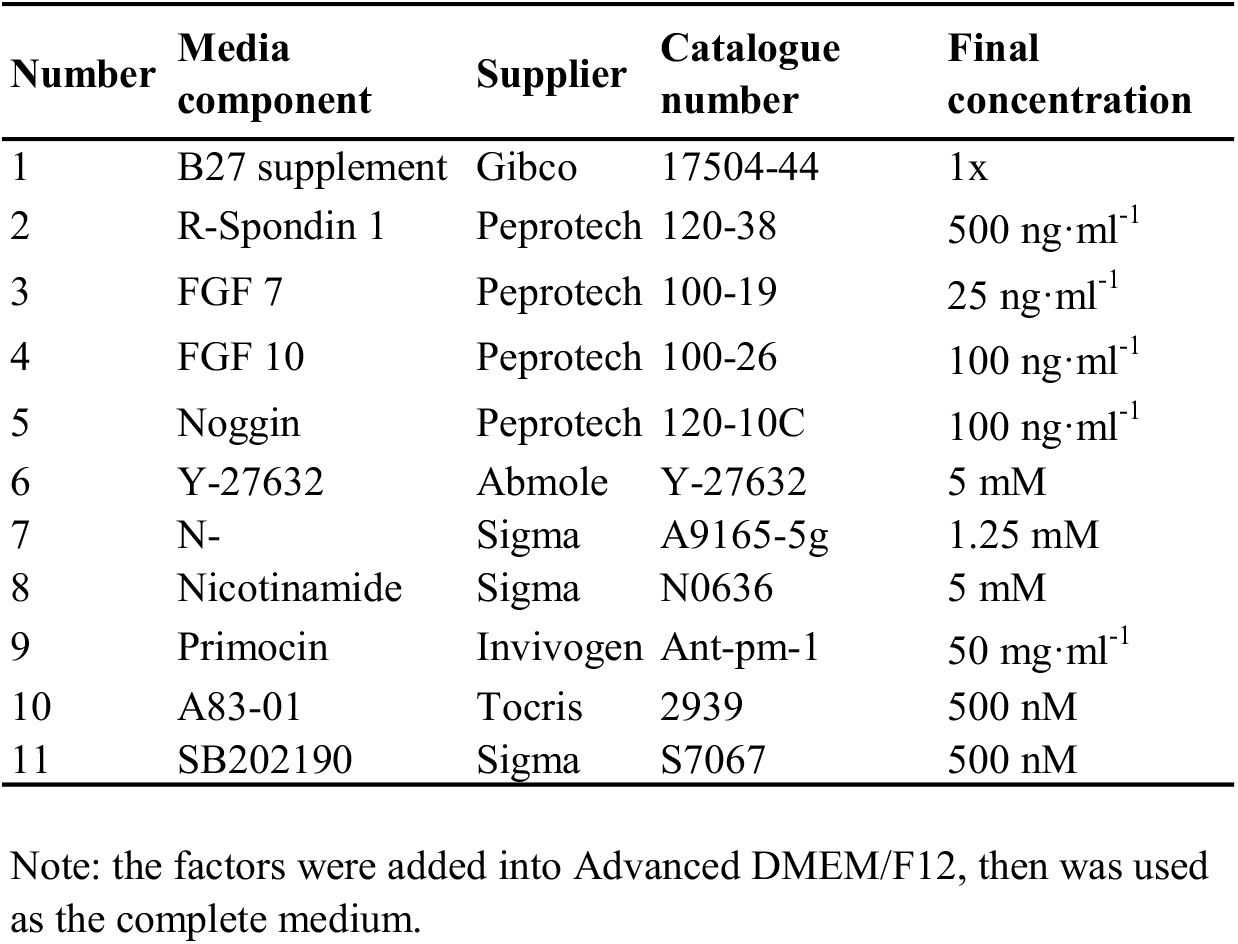
Lung Organoid Medium Recip

